# Stochastic gene expression influences the selection of antibiotic resistance mutations

**DOI:** 10.1101/563015

**Authors:** Lei Sun, Peter Ashcroft, Martin Ackermann, Sebastian Bonhoeffer

**Affiliations:** Institute of Integrative Biology, ETH Zürich, Zürich, Switzerland; Dept. of Systems Biology, Harvard Medical School, Boston, MA, USA; Institute of Biogeochemistry and Pollutant Dynamics, ETH Zürich, Zürich, Switzerland; Department of Environmental Microbiology, EAWAG, Swiss Federal Institute of Aquatic Science and Technology, Dübendorf, Switzerland

## Abstract

Bacteria can resist antibiotics by expressing enzymes that remove or deactivate drug molecules. Here, we study the effects of gene expression stochasticity on efflux and enzymatic resistance. We construct an agent-based model that stochastically simulates multiple biochemical processes in the cell and we observe the growth and survival dynamics of the cell population. Resistance-enhancing mutations are introduced by varying parameters that control the enzyme expression or efficacy. We find that stochastic gene expression can cause complex dynamics in terms of survival and extinction for these mutants. Regulatory mutations, which augment the frequency and duration of resistance gene transcription, can provide limited resistance by increasing mean expression. Structural mutations, which modify the enzyme or efflux efficacy, provide most resistance by improving the binding affinity of the resistance protein to the antibiotic; increasing the enzyme’s catalytic rate alone does not contribute to resistance to ciprofloxacin, but may contribute if drug binding isn’t rate limiting. Overall, we identify conditions where regulatory mutations are selected over structural mutations, and vice versa. Our findings show that stochastic gene expression is a key factor underlying efflux and enzymatic resistances and should be taken into consideration in future antibiotic research.

## 1 Introduction

Efflux pumps and drug-inactivating enzymes allow bacterial cells to evade the damage caused by antibiotic drugs. Bacterial pathogens with efflux and enzymatic resistances are ubiquitous and often are serious public health concerns (Li et al., 2015; World Health Organization, 2017). For instance, several gram-negative bacteria species with carbapenemase and ESBL-activities are listed as critical priority pathogens for research and development of new antibiotics by the WHO (World Health Organization, 2017).

Studies to quantify antibiotic resistance are routinely performed at the bacterial population level, i.e. through protocols such as minimum inhibitory concentration (MIC) or time-kill curves. However, the action of the drug occurs within the cell on the molecular level. Bridging these different scales remains a challenge.

The action of antibiotic drugs usually involves small numbers of molecules and binding sites, such that intrinsic stochasticity could have a significant effect on the dynamics. This noise impacts how we interpret bacterial assays that extract population-averaged behaviours, which normally ignore cell-to-cell heterogeneity. A recent study has suggested that many experimental phenomena, such as post-antibiotic effects, can be explained by these noisy within-cell dynamics (Abel Zur Wiesch et al., 2015).

Another source of randomness is in the mechanism of gene expression (Paulsson, 2005). Transcription typically occurs in bursts, a phenomenon often described by the so-called two-state or telegraph model of gene expression (Paulsson, 2005; Kumar et al., 2015; Lionnet and Singer, 2012). In this model, gene transcription switches stochastically between an active state with a constant rate of mRNA production, and an inactive state without transcription. Although certain genes may have complex transcription regulations, the two-state model is widely used when modelling both prokaryotes and eukaryotes due to its simplicity (Paulsson, 2005). The duration of active and inactive states can range from minutes to multiple cell generations (Lionnet and Singer, 2012; Hammar et al., 2014; So et al., 2011). The intrinsic stochasticity of gene expression is often the dominant source of randomness for gene-product numbers, and the separation of noise intensities has been used to perform mathematical analysis of gene-expression systems (Lin and Galla, 2016). While the expression level of resistance enzymes or efflux pumps generally correlates with the level of observed phenotypic resistance (Zwart et al., 2018), the effects of stochasticity on resistance remain unexplored.

Enzymatic resistances can be enhanced by regulatory or structural gain-of-function mutations, affecting the gene expression or the enzyme efficacy, respectively (Händel et al., 2014; Berrazeg et al., 2015; Yang et al., 2003; Blair et al., 2015). Both regulatory and structural mutations have been described for a number of antibiotic resistances (Händel et al., 2014; Toprak et al., 2012; Berrazeg et al., 2015). However, the selective conditions that favour one type of mutation over the other have not been fully explored.

Here, we provide a computational model that investigates the effects of stochastic gene expression on resistance and resistance evolution. Our model describes the dynamics of within-cell processes of the drug-target-efflux system and accounts for stochasticity in transcription and translation, as well as drug diffusion, binding, and removal. Using this model, we consider regulatory and structural resistance mutations to study their behaviour on the molecular, cellular, and population levels. Our goal is to explore how stochastic gene expression influences the survival and extinction of different types of resistance mutations under antibiotic treatments.

## 2 Model

### 2.1 Overview

We model a population of elongating and dividing cells in a boundless environment with an antimicrobial drug. The drug molecules can diffuse across cell membranes, and they influence the cell’s death rate by binding to drug-specific targets. A drug efflux system, which is subject to stochastic gene expression, removes drug molecules from the cell. We focus on mutations that affect this system, either through regulatory effects or changes to the enzymatic efficacy.

Each cell is modelled as an independent agent, such that there are no between-cell interactions. To uphold independence, we assume a constant concentration of drug outside the cell. Spatial effects are ignored. The within-cell model takes the following processes into account: cell elongation and division; intracellular production of drug-target protein; entry of drug molecules into the cell; the interaction of target proteins with drugs and subsequent cell death due to antibiotics; and the transcription, translation and enzymatic activity of the resistance-mediating proteins. From here on we refer to the resistance-mediating protein as efflux protein, as it can remove drug molecules from the cell either as an efflux pump or drug-inactivation enzyme.

Each cell is described by seven discrete variables (shown in Table 1), which are updated stochastically using the adaptive tau-leaping Gillespie algorithm as implemented in the `adaptivetau` package in `R` (Cao et al., 2007). This is an approximation of the full stochastic simulation algorithm (SSA) (Gillespie, 1977). The interactions between the within-cell variables can be seen in the model schematic in Fig. 1 and are described in detail below. In addition, we model the elongation of the cell deterministically. All cell parameter values are reported in Supplementary Table S2.

**Table 1:**
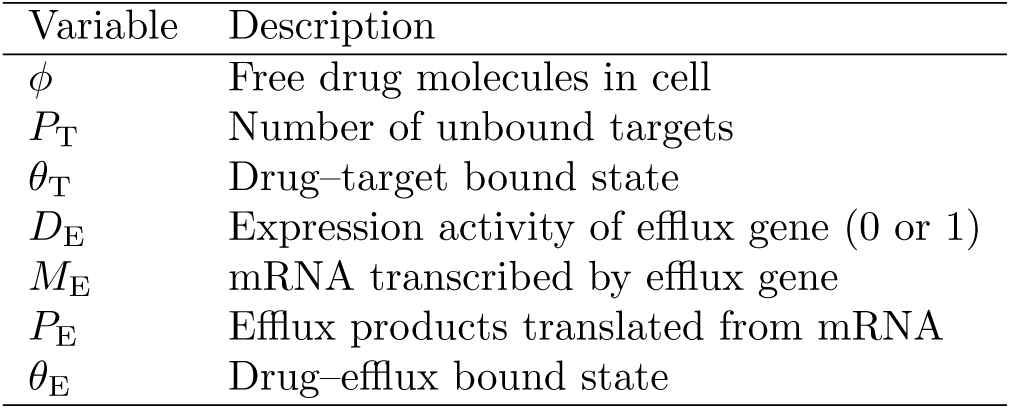
Within-cell variables

| Table 1: Within-cell variables |  |
| --- | --- |
| Variable | Description |
| $\phi$ | Free drug molecules in cell |
| $P_T$ | Number of unbound targets |
| $\theta_T$ | Drug-target bound state |
| $D_E$ | Expression activity of efflux gene (0 or 1) |
| $M_E$ | mRNA transcribed by efflux gene |
| $P_E$ | Efflux products translated from mRNA |
| $\theta_E$ | Drug-efflux bound state |

**Figure 1:**
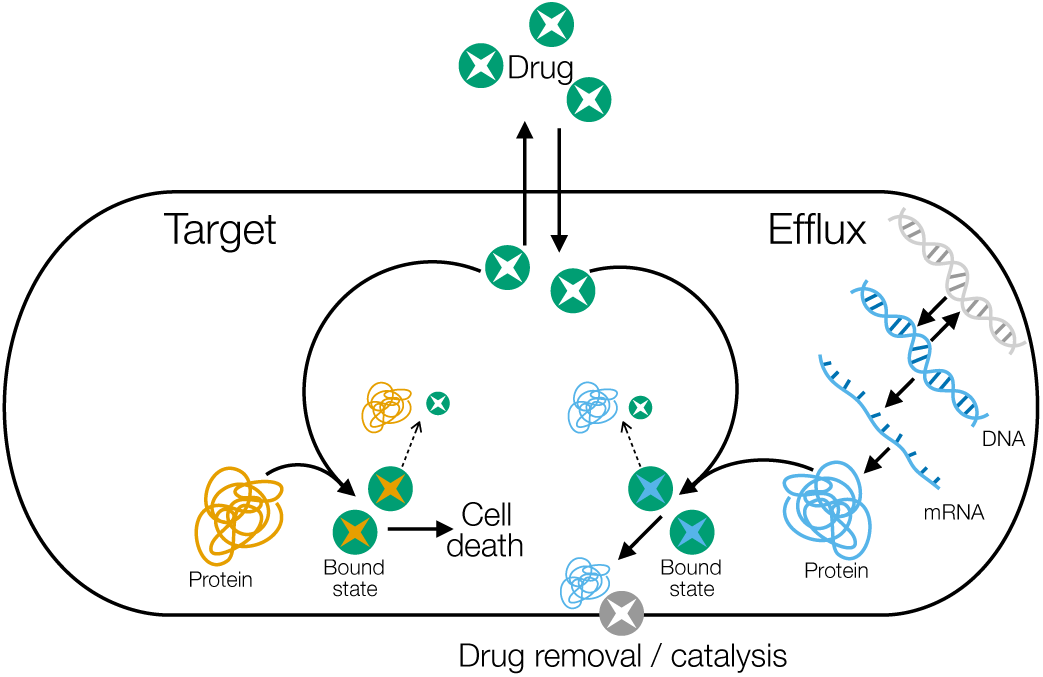
Model schematic. Each arrow corresponds to a discrete reaction, as described below. Dashed arrows indicate that drug–protein bound states may dissociate (depending on the drug used). Efflux proteins are translated from mRNA, which in turn is transcribed from DNA. The DNA can be active (blue) or inactive (grey), and can switch between these two states. Not shown is the constant production of target proteins, which we assume are constitutively expressed.

### 2.2 Cell physiology and division

The cells we model here *are Escherichia coli*. We assume that the cells are cylindrical with cross-sectional diameter *d*. A newly-divided cell has length *ℓ*_0_. The cell elongates exponentially until it reaches twice its length at birth (Campos et al., 2014), which is achieved in time interval *t*_*G*_. At a time *t* after birth, the cell length (*ℓ*), surface area (*A*), and volume (*V*) satisfy

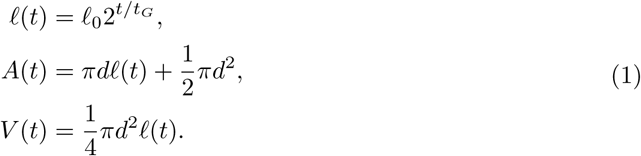

For each cell, a small amount of zero-expectation Gaussian noise (standard deviation = 5%) is added to the mean generation time (*t*_*g*_ → 𝒩 [*t*_*G*_, (0.05*t*_*G*_)^2^]) to desynchronise cell divisions and to increase cell-to-cell heterogeneity. Division happens instantaneously once the cell reaches twice its birth-length, and produces two equal-length daughter cells. All variables in Table 1 are divided binomially between the two daughter cells upon cell division, with the exception of the DNA activity *D*_E_, which is directly inherited by both daughters.

### 2.3 Target production

The target proteins usually fulfil essential functions in the cell, such as ribosomal subunits, RNA polymerase, DNA gyrase, and cell-wall components (Andersson and Hughes, 2010). The numbers per cell of several targets have been well-characterised and used for computational studies (Abel Zur Wiesch et al., 2015). The intracellular concentrations of these essential proteins may impact the cell’s growth, for instance as in the case with ribosomes (Greulich et al., 2015). Therefore, we assume that these target proteins are subject to various intracellular regulations that maintain a stable and optimal concentration within the cell. This assumption is consistent with previous findings showing essential genes have low protein-expression noise (Silander et al., 2012). In the model this is achieved through constant production of targets over the duration of the cell’s life. The production rate Γ is chosen such that, on average, the number of targets doubles in a generation. I.e., 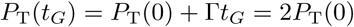, such that

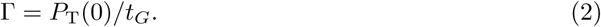

As the half-life of bacterial proteins is usually on the scale of multiple hours (Larrabee et al., 1980), we neglect protein degradation as it is relatively insignificant to dilution-by-cell-division.

### 2.4 Efflux production

The transcription and translation of efflux proteins are explicitly modelled: we expect the expression of these non-essential genes to be more stochastic than that of the drug targets. The gene activity switches on(off) with a fixed rate Γ_On_(Γ_Off_), and therefore follows the telegraph process of gene expression (van Kampen, 2007). Once the gene is activated (*D*_E_ = 1), mRNA is transcribed with a constant rate *τ*. The mRNA molecules degrade with rate *γ*. On average, *b* proteins are translated during the lifetime of each mRNA molecule. As above, we assume that these proteins do not decay but are diluted only by cell division. Given that these processes are intrinsically stochastic, a brief period of gene activation may not always produce efflux mRNA and/or protein.

### 2.5 Drug diffusion, interaction, and efflux

The antibiotic drug outside of the cells is kept at a constant concentration *c*_out_. The drug molecules diffuse through the cell envelope bi-directionally with a fixed diffusion rate *σ*. The overall diffusion rate is proportional to the surface area of the cell, *A*(*t*). We stochastically model Fick’s law of diffusion for the bi-directional movement, such that transport is driven by a concentration gradient.

Once a drug molecule has entered the cell, it can either diffuse out, bind to its specific target, or be captured by an efflux protein. These binding events result in the formation of a drug–protein bound state. Binding occurs with the forward rate constant *k*_*f*_, reflecting the binding affinity of target and efflux proteins. The actual binding rate also depends on the concentration of the drug and the proteins that interact inside the cell. Drug-binding to target and efflux are both considered reversible unless stated otherwise, and the bound states dissociate with backwards rate *k*_*b*_. This event releases both the drug molecule and the protein into the cell. To compete efficiently with targets, efflux proteins should have at least the same or higher drug-binding rates than target proteins. Therefore we set the basal efflux binding rate to be the same as that of targets, making them indistinguishable to the drug. The efflux protein has the capacity to remove the drug molecule once bound, through catalysis or transport, and this happens with rate *k*_cat_.

All of the above reactions are summarised in Table 2.

**Table 2:** Within-cell reactions

| Table 2: Within-cell reactions |  |  |
| --- | --- | --- |
| Reaction | Rate | Description |
| $\emptyset \rightarrow P_T$ | $\Gamma$ | Constant production of targets |
| $\emptyset \rightarrow D_E$ | $\Gamma_{\text{On}}(1 - D_E)$ | Switch on efflux gene |
| $D_E \rightarrow \emptyset$ | $\Gamma_{\text{Off}}D_E$ | Switch off efflux gene |
| $D_E \rightarrow D_E + M_E$ | $\tau D_E$ | Transcription of efflux mRNA |
| $M_E \rightarrow \emptyset$ | $\gamma M_E$ | Decay of efflux mRNA |
| $M_E \rightarrow M_E + P_E$ | $\gamma b M_E$ | Translation of efflux product |
| $\emptyset \rightarrow \phi$ | $\sigma A(t)c_{\text{out}}$ | Influx of free drug into cell |
| $\phi \rightarrow \emptyset$ | $\sigma A(t)\phi/V(t)$ | Loss of free drug from cell |
| $\phi + P_T \rightarrow \theta_T$ | $k_f P_T \phi/V(t)$ | Drug-target binding |
| $\theta_T \rightarrow \phi + P_T$ | $k_b \theta_T$ | Dissociation of drug-target bound state |
| $\phi + P_E \rightarrow \theta_E$ | $k_f P_E \phi/V(t)$ | Drug-efflux binding |
| $\theta_E \rightarrow \phi + P_E$ | $k_b \theta_E$ | Dissociation of drug-efflux bound state |
| $\theta_E \rightarrow P_E$ | $k_{\text{cat}} \theta_E$ | Catalysis/removal of drug |

### 2.6 Cell death and MIC fraction

Here we consider bactericidal drugs that introduce a drug-dependent death rate for each cell. Therefore, the presence of drug has no impact on the cell generation time. We combine two previously published models to implement a realistic death mechanism: the drug-induced death rate, log(10)*δ*(*ρ*), is an increasing function of the fraction of bound targets, *ρ* (Abel Zur Wiesch et al., 2015), but has a maximum value (Regoes et al., 2004). Under these conditions, the number of cells n follows 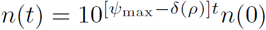, where *Ψ*_max_ is the population growth rate in the absence of antibiotic. The fraction of bound targets, *ρ*, also correlates with the concentration of bound targets.

The death rate should satisfy the following criteria:

1. *δ*(0) = 0: In the absence of antibiotics the death rate is zero;
2. *δ*(*ρ*_MIC_) = *Ψ*_max_: When the external drug concentration is 1×MIC – which is the minimum concentration at which no bacterial growth is observed, the fraction of bound targets is given as *ρ*_MIC_. The death rate at this value should equal the maximum growth rate in the absence of antibiotic (*Ψ*_max_), such that the net growth rate is zero;
3. *δ*(1) = *Ψ*_max_ − *Ψ*_min_: When all targets are bound, the death rate is maximal. The net growth rate is then *Ψ*_min_., which is the minimum measured population growth rate.

In our model, the death rate takes the sigmoidal form as in Regoes et al. (2004),

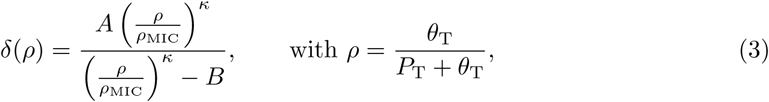

where the constants *A* and *B* are chosen such that the above criteria are satisfied, i.e.,

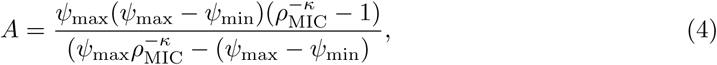

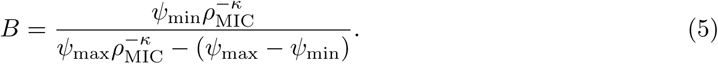

The values of *Ψ*_max_ and *Ψ*_min_ have been determined for multiple antibiotic compounds (Regoes et al., 2004). Finally, the shape parameter *κ* in Eq. (3) determines the steepness of the sigmoidal response. For *κ* → ∞, we recover the step function where death only occurs if the fraction of bound targets is greater than a threshold (in this case, the threshold would be *ρ*_MIC_).

Through Eq. (3) we can relate the within-cell parameter *ρ*_MIC_ to the externally-measured (population-level) MIC of the drug. Empirically, MIC is determined as the lowest drug concentration to prevent a small bacterial inoculum from proliferating overnight into visible density, sometimes quantified specifically as > 90% reduction in growth rate compared to a drug-free control (Arthington-Skaggs et al., 2002; Tunney et al., 2004; Toprak et al., 2012). Alternatively, one could define the MIC as the concentration of drug at which the net growth rate is zero after a given time period; this is referred to as zMIC by Regoes et al. (2004). In the Supplementary Material, we measure these quantities in simulations and compare them to different lineage survival probability thresholds following overnight experiments, such as IC50 (50% lineage survival), IC90 (10% lineage survival), and IC99 (1% lineage survival) (Supplementary Fig. S1). The reason for using lineage survival probability is its computational efficiency compared to computing population growth rates. We find that IC90 is a suitable measure of MIC which correlates well with the zMIC drug concentration.

We determined values of *ρ*_MIC_ by simulating lineages emerging from individual wildtype cells that grow for 20 hours in a constant environment with *c*_out_ = 1×MIC. This time interval is comparable to standard overnight MIC-experiments (Regoes et al., 2004). We then screened for *ρ*_MIC_, defined as the highest value of bound target fraction where more than 90% of the progenitor cells and their lineages become extinct. This screening process is highlighted in Supplementary Fig. S2 for the drug ciprofloxacin, from which we find an MIC-fraction of 8.1%. Although this number may seem low, it could potentially be explained by the drug mechanism: ciprofloxacin binds to DNA-bound gyrases and fragments the bacterial chromosome via DNA doublestrand-breaks (DSBs) (Kampranis and Maxwell, 1998; Tamayo et al., 2009). Given that each *E. coli* cell has ∼ 300 DNA-bound gyrases (Chong et al., 2014), this MIC-fraction suggests that a cell is likely to die when its chromosome is fragmented by over 20 simultaneous DSBs, which seems plausible. The drug-specific parameters are listed in Supplementary Table S3, and we repeat this screening procedure for rifampicin (Supplementary Fig. S3).

### 2.7 Simulation algorithm

To simulate a population of cells over multiple generations, we use the following algorithm:

1. Assign each initial cell a unique set of parameter values;
2. Identify the cell with the earliest birth time;
3. Run the adaptive tau-leaping simulation algorithm for this cell until the cell dies, or it reaches its predefined generation time;
4. If the cell survived, perform cell division to create two new daughters;
5. Repeat from 2. until the experiment ends, no cells remain, or the number of cells is large enough that we can assume survival of the population for the duration of the experiment (200 unless otherwise stated).

We also describe the within-cell behaviour by a system of ODEs, as shown in the Supplementary Material. We refer to these equations as the mean-field solution, as it does not take into account any stochasticity, and instead reflects the average dynamics of an infinite number of cells.

### 2.8 Model behaviour

To test the validity of our model, we checked the distribution of target and efflux molecules across a large ensemble of wildtype cells growing in the absence of antibiotics (Supplementary Fig. S4). We found that the frequency of cells with active efflux DNA agrees with the prediction of the telegraph process (∼ 2.5%), while the average number of efflux mRNA (∼ 0.04) and efflux proteins (∼ 4) per cell are within experimentally reported ranges (Moran et al., 2013; Cai et al., 2006; Taniguchi et al., 2010). The efflux protein copy number also follows a negative-binomial distribution, as predicted by the theory of their bursty dynamics (Paulsson and Ehrenberg, 2000). Finally, the number of target proteins per cell is normally distributed, which is expected given its constant production rate.

In Fig. 2 we show an example trajectory of the intracellular model variables in the absence and presence of an antibiotic (ciprofloxacin). Starting from a single wildtype cell, we track only one daughter cell after each division. After six generations we instate an external drug concentration, such that the drug can freely diffuse into the cell (Fig. 2A). The mean-field equations closely approximate the number of constantly-produced intracellular drug targets (Fig. 2B). The number and fraction of drug-bound targets is also well approximated in the mean-field limit (Fig. 2C,D). However, the bursty dynamics of the efflux system are not well captured by the deterministic approximation (Fig. 2E–H). Here a piecewise deterministic Markov process, which accounts for the gene-expression noise, would be a more appropriate approximation for this system (Lin and Galla, 2016).

**Figure 2:**
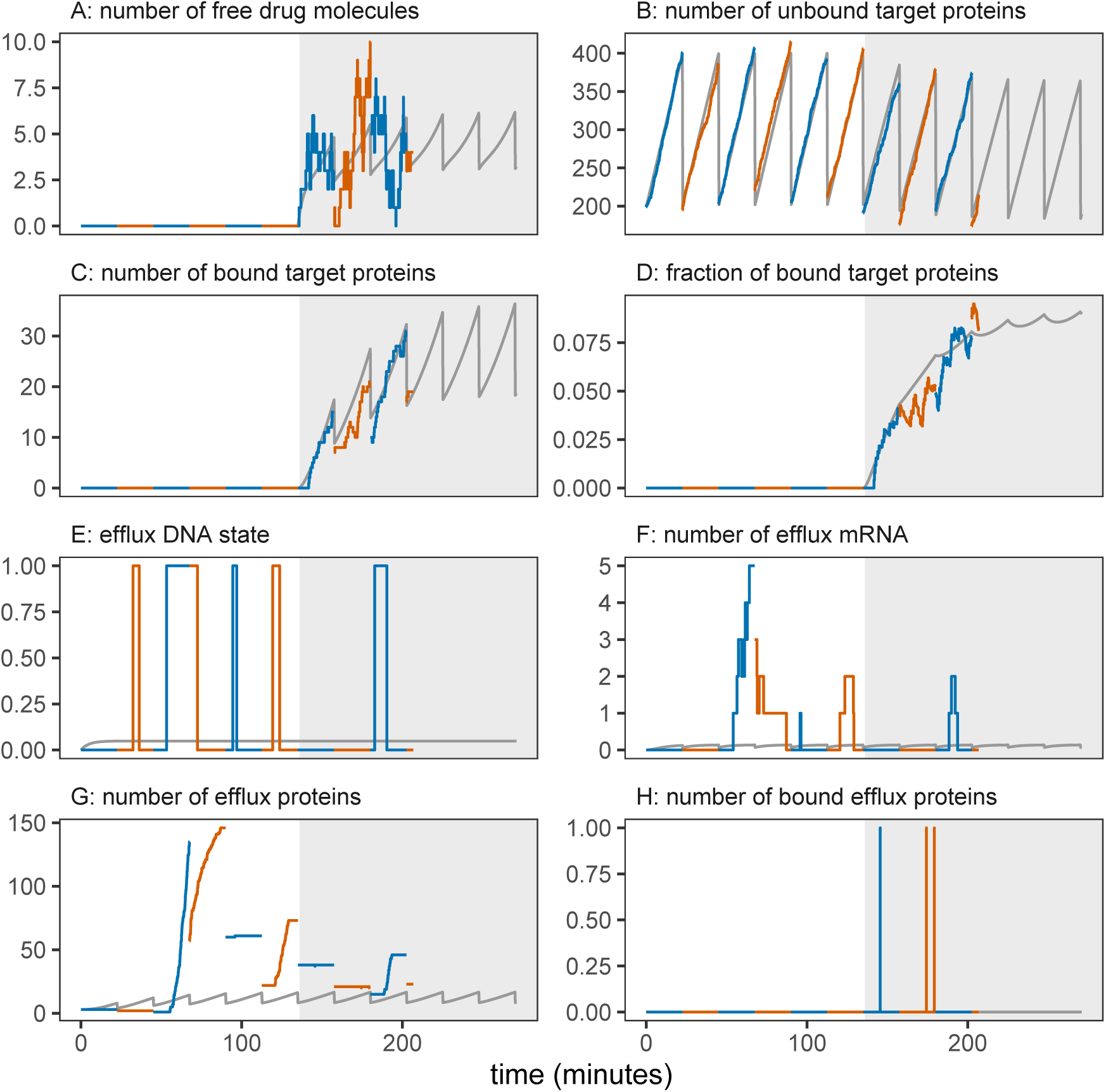
Intracellular variables in the absence (white background) and presence (grey background) of ciprofloxacin. Cell generations are distinguished by alternating line colors. Solid grey lines show the expected values based on mean-field solutions [Supplementary Material Eq. (S1)]. The external drug concentration is set to zero for the first six generations, and then to *c*_out_ = 1 × MIC. The tracked cell lineage died after ∼ 200 minutes. The panels show: A) number of free intracellular drug molecules; B,C) number of free and drug-bound target proteins, respectively; D) fraction of target proteins that are bound by the drug; E) stochastic expression of the efflux gene; F) number of efflux mRNA in the cell; G,H) number of free and drug-bound efflux proteins, respectively. Model parameters are listed in Supplementary Tables S2 and S3, but we double the DNA activation rate Γ _On_ to more clearly illustrate the stochastic gene expression dynamics in a shorter time window.

Although cells could accumulate multiple efflux proteins during the period of gene activity (Fig. 2G), we did not observe two or more drug-bound efflux proteins at the same time (Fig. 2H). This results from two factors: the binding rate of drug is slow enough to disallow near-simultaneous formation of multiple drug–efflux bound states, while the catalysis rate is high enough that any efflux-bound drug molecule is quickly catalysed. Put together, it shows that here the rate-limiting factor for removing drugs by efflux is not the catalysis step, but the drug-binding process (at least for this set of parameters).

### 2.9 Mutations

We consider seven classes of cells throughout this study: knockout, wildtype, three types of regulatory mutants and two types of structural mutants. Knockout (KO) cells lack efflux protein production. Wildtype (WT) cells have the basal level of efflux production. The effects of a mutation are characterised by a multiplicative parameter *µ* > 1. The three types of regulatory mutants we consider are: REG-ON, which has an increased rate of gene activation (Γ_On_ → *µ*Γ_On_); REG-OFF, which has longer periods of active transcription by reducing the inactivation rate (Γ_Off_ → Γ_Off_ */µ*); and REG-BURST, which increases the translation rate or protein burst size from each translational event (*b* → *µb*). The two types of structural mutants include STRUCT-BIND with improved binding affinity (*k*_*f*_ → *µk*_*f*_) to drug molecules and STRUCT-CAT with improved catalytic rates to break down drug molecules (*k*_cat_ → *µk*_cat_). The mutant cells also carry a cost, 0 < *ν* ≪ 1, which is associated with their modified function. We assume this cost affects all metabolic processes in the cell, leading to a longer generation time, as well as slower translation. We therefore implement Γ → (1 − *ν*)Γ, *b* → (1 − *ν*)*b*, and ⟨*t*_*G*_⟩ → ⟨*t*_*G*_⟩ */*(1 − *ν*). The KO cell type carries a negative cost, as we assume it grows slightly faster for lacking efflux protein production completely.

## 3 Results

### 3.1 Lineage survival

We first simulate the survival probability of cell lineages when faced with a constant concentration of ciprofloxacin for 20 hours. As well as considering different mutant classes, we also vary the mutant effect parameter *µ*. KO and WT classes are included as controls.

Four classes of mutant show distinct survival probability profiles (Fig. 3). REG-BURST mutants share an almost-identical profile to STRUCT-BIND, therefore the results for this class are relegated to the Supplementary Material (Supplementary Fig. S5). REG-ON mutants are more likely to survive at low drug concentrations (near the WT MIC) than any other mutant class, but their survival probabilities decline sharply with increasing drug concentrations (Fig. 3A). Death is more prevalent at low concentrations for REG-OFF mutants when compared to REG-ON, but the decrease in survival probability with concentration is slower (Fig. 3B). Ultimately, REG-ON and REG-OFF mutants are eliminated at similar concentrations. STRUCT-BIND mutants (and also REG-BURST) have much lower survival probabilities than regulatory mutants at low drug concentrations (Fig. 3C). However, a small fraction of STRUCT-BIND mutants can survive high drug concentrations beyond what the regulatory mutants can survive. STRUCTCAT mutants, on the other hand, do not show any noticeable improvement in survival probability compared to WT or KO (Fig. 3D).

**Figure 3:**
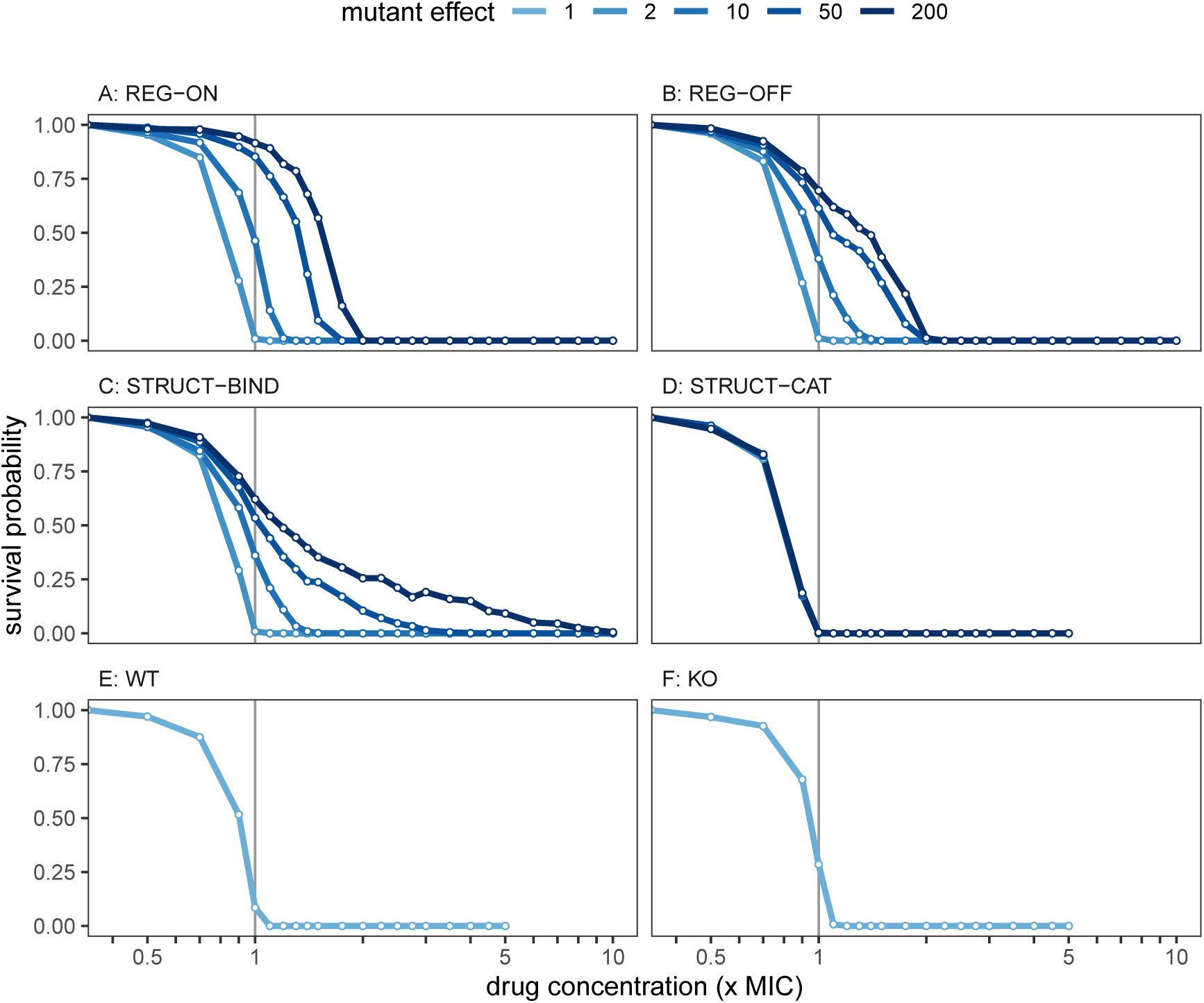
The survival probability of the different mutant classes across increasing concentrations of ciprofloxacin. Drug concentration is measured relative to the wildtype MIC. Each data point is the mean of 1,000 individual simulations, with each simulation starting from one individual cell of the respective mutant class. Each simulation ends when either the population size reaches 200 cells (at which point extinction is extremely unlikely), all cells die, or we reach the end of the experiment at *t* = 1, 200 minutes. Survival probability indicates the fraction of simulations with more than zero live cells.

From Fig. 3 we can interpolate IC90 values (external drug concentration at which lineage survival is 10%) for each cell type. Both regulatory mutations (REG-ON and REG-OFF) are characterised by successive increase in their IC90 values with increasing mutant effect. At *µ* = 200, the regulatory mutations have an IC90 value that is increased by almost two-fold, which is comparable to the measured three-fold increase in MIC for a regulatory mutation in *acrR* (Marcusson et al., 2009). STRUCT-BIND mutants show a small increase in IC50, but significant increase in IC90 with increased binding rate, reaching 4.5-fold increase at *µ* = 200. STRUCTCAT has slightly lower IC values than WT, due to fitness cost of resistance and no obvious benefit of increasing the rate of catalysis. Conversely, KO has slightly higher IC values than WT due to its small fitness advantage. The IC90 values, as well as IC50 values, for all cells and mutant effect values are reported in Supplementary Table S4.

For the cell lineages that became extinct in Fig. 3, we looked at their extinction times (Supplementary Fig. S6). Like the WT and KO cell types, the structural mutants and REGBURST have their peak mean extinction time near the wildtype MIC. If extinction occurs at sub-MIC concentrations (which is a rare event, as shown in Fig. 3), it is most likely to happen at the beginning of the simulation when cell numbers are small. At 1×MIC, the extinction time diverges as death rate equals growth rate at this drug concentration (so that the expected lineage lifetime becomes infinite). As the drug concentration increases beyond the MIC, more drug diffuses into the cells and the death rate increases, leading to faster extinction of the cell lineages. Compared to WT and KO, both REG-ON and REG-OFF mutants show a shift of the peak extinction time towards higher concentrations with larger mutational effects. This corresponds to the shift in their respective IC90 values. Finally, we note that the STRUCTBIND lineages of cells show a broad distribution (large variance) of extinction times at high drug doses.

Combining the data on survival probability and extinction times, our results suggest that REG-ON mutations provide intermediate but homogeneous resistance. This is characterised by the high survival probability at low drug concentrations, which declines quickly as the drug concentrations increase until all cells eventually die. Resistance resulting from a REG-OFF mutation, however, seems heterogeneous. This is characterised by prominent deaths even at low concentrations but with some cell lines surviving through intermediate drug doses. The heterogeneity is even more pronounced in the STRUCT-BIND mutants: the low extinction times show that some cells are dying early, but the continued survival through intermediate and high doses suggest that a fraction of the cells is highly resistant. For STRUCT-BIND, the extreme resistance heterogeneity should be the combined result of low but highly stochastic gene expression and elevated binding rates of the resistance enzyme, such that the few cells with copies of the improved efflux protein are highly protected.

### 3.2 Growth rates

An alternative measure of mutant performance is through their population-level growth rates across different drug concentrations. As well as measuring the level of resistance conferred, these results can be directly compared with empirical observations. Following the protocol of Regoes et al. (2004), we simulated populations of cells for a short duration and counted the number of live cells at 10-minute intervals, before extracting the net population growth rate (Fig. 4).

**Figure 4:**
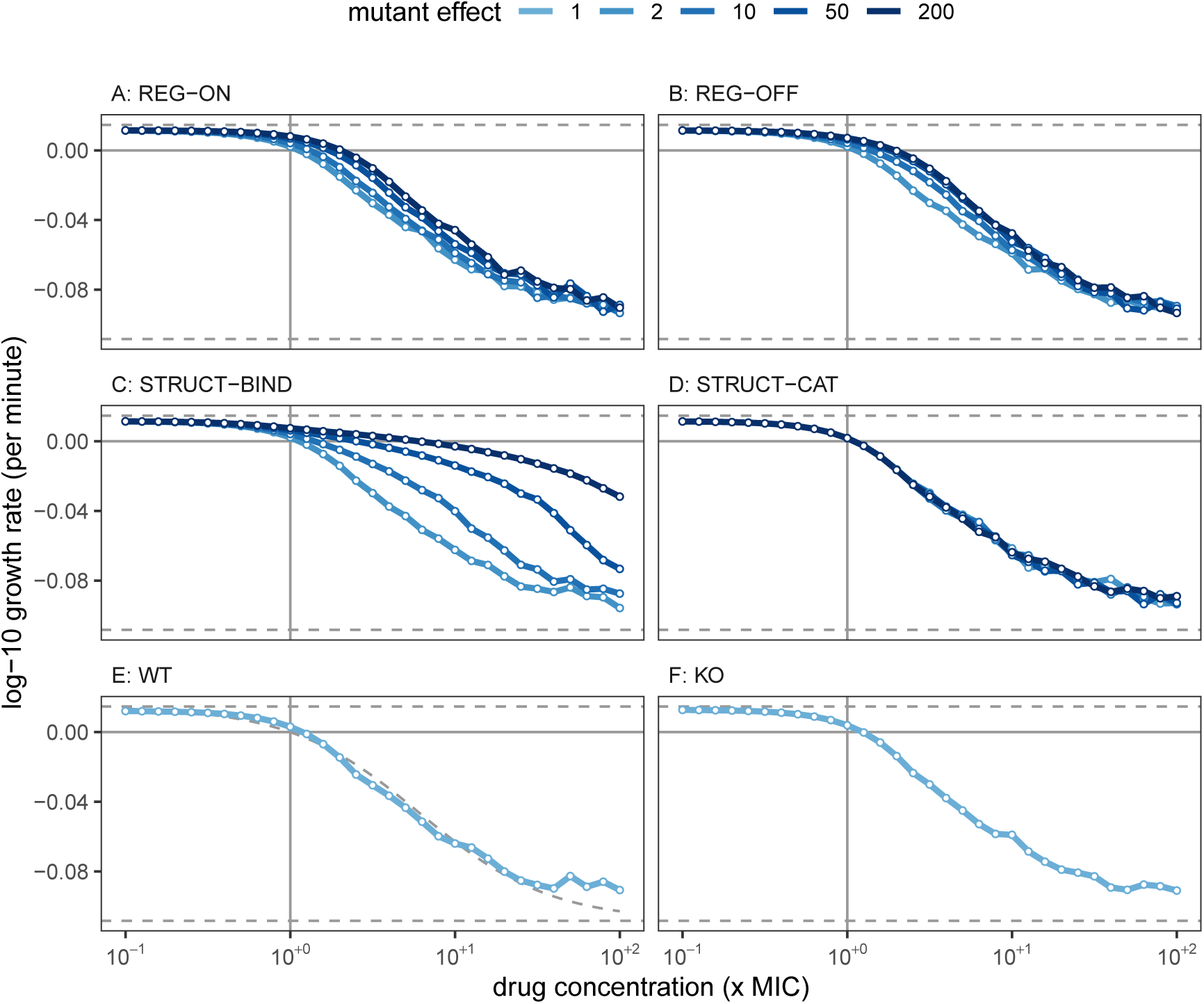
Population growth rates during three hours of antibiotic exposure. For each mutant class and effect level, we simulated a population of up to 100,000 cells, and recorded the number of live cells at 10 minute intervals. From the corresponding time-kill curves (shown in Supplementary Fig. S7), we extracted the net growth rate as the linear regression coefficient between the base-10 logarithm of cell number and the sample time, as described by Regoes et al. (2004). Horizontal dashed lines are the maximum and minimum growth rates, *Ψ*_max_ and *Ψ*_min_. The dashed curve in panel E is the dose-response curve from Regoes et al. (2004).

Both regulatory mutants show similar dose-response profiles, which gives no indication of the differences in their lineage survival probability. STRUCT-BIND mutants, however, generally maintain a higher net growth rate under high drug pressure when compared to the other cells types. This is in agreement with Fig. 3, where a few cell lineages are able to proliferate at these high treatment intensities. STRUCT-CAT, WT, and KO cells show the classical sigmoidal dose response, and the WT dose-response curve agrees well with the data reported by Regoes et al. (2004) (Fig. 4E).

### 3.3 Heterogeneity and resistance

The number of efflux proteins per cell in REG-ON and REG-OFF mutants are nearly identical in mean for a given mutation effect *µ* (Fig. 5). However, these mutant classes have very different distributions of efflux protein number across the population. For REG-ON mutants the protein number distribution is unimodal such that cells are more homogeneous; with frequent transcription bursts (e.g. at *µ* = 50 and 200) all cells produce some efflux proteins. REG-OFF mutants, in contrast, display a bimodal distribution of protein number: A large fraction of cells within the population do not produce any efflux protein, while some produce more than the corresponding REG-ON mutant. This is the cause of REG-OFF cell death even at low drug concentrations, but with some survival at intermediate doses.

**Figure 5:**
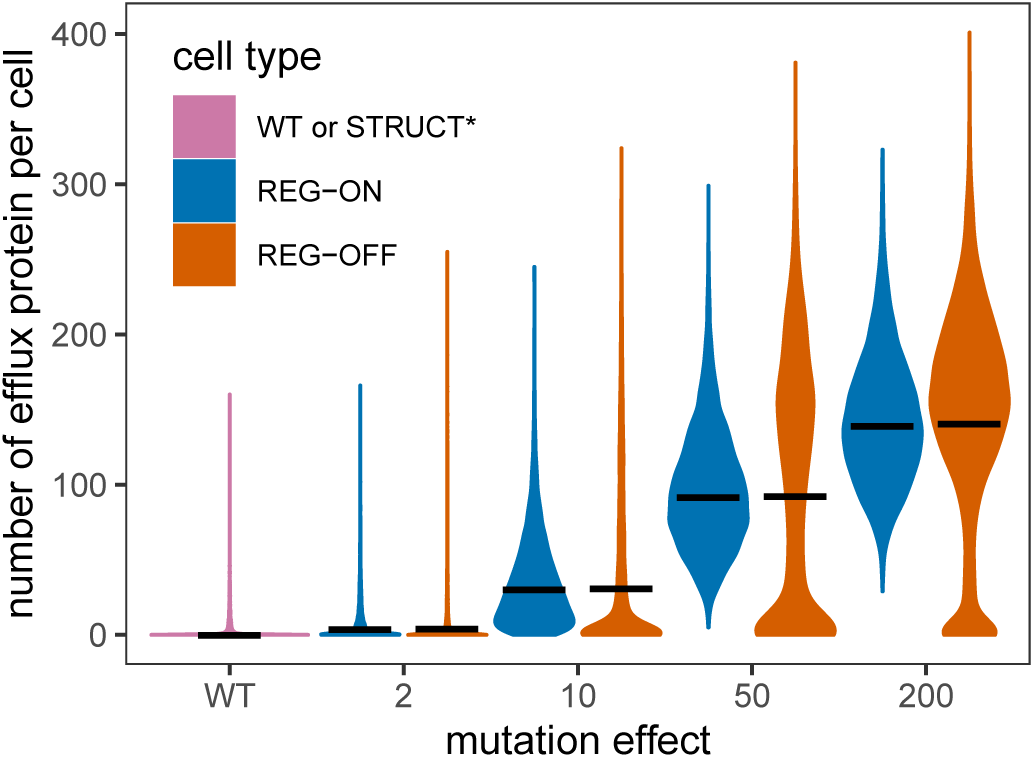
The distribution of efflux protein copy number per cell for regulatory mutants, across a range of mutant effects *µ*. Wildtype (WT) cells are included for comparison, and the expression of structural mutants is identical to WT. The mean protein copy numbers per cell are indicated by solid black lines. A population of 10,000 individual cells was simulated up to *t* = 1, 200 minutes in the absence of drug, tracking only a single cell following each division event. Numbers are recorded immediately after cell division.

Overall, regulatory mutations only provide limited resistance. For REG-ON and REG-OFF with 200× mutational effects, transcription is active for ∼ 80% of the time. Therefore, resistance cannot be improved significantly by augmenting the transcription dynamics beyond this level. Further improvements in resistance then rely on increasing the translation rate (resulting in more proteins per mRNA) or by improving the efficacy of the efflux protein itself. This is consistent with previous findings showing that resistance provided by overexpression is limited (Toprak et al., 2012; Yang et al., 2003).

For STRUCT-BIND mutants, the efflux protein number distribution is very similar to that of the wildtype as there are no changes in expression (apart from the small cost of carrying the mutation). Therefore, the majority of STRUCT-BIND cells in a population contain no efflux protein. However, those cells that do contain efflux proteins have a high level of resistance, due to the high efficacy of these proteins. This is the source of heterogeneity in the STRUCT-BIND mutant population.

To investigate this further, we consider the survival for a STRUCT-BIND mutant, but with different levels of gene expression stochasticity (Fig. 6). Concretely, we increase the frequency of gene activation (Γ_On_ → *α*Γ_On_ for *α* > 1), and correspondingly decrease the efflux protein translation rate *γb* → *γb/α*. In this way, we maintain the mean number of efflux proteins per cell (Fig. 6A) but alter the expression noise.

**Figure 6:**
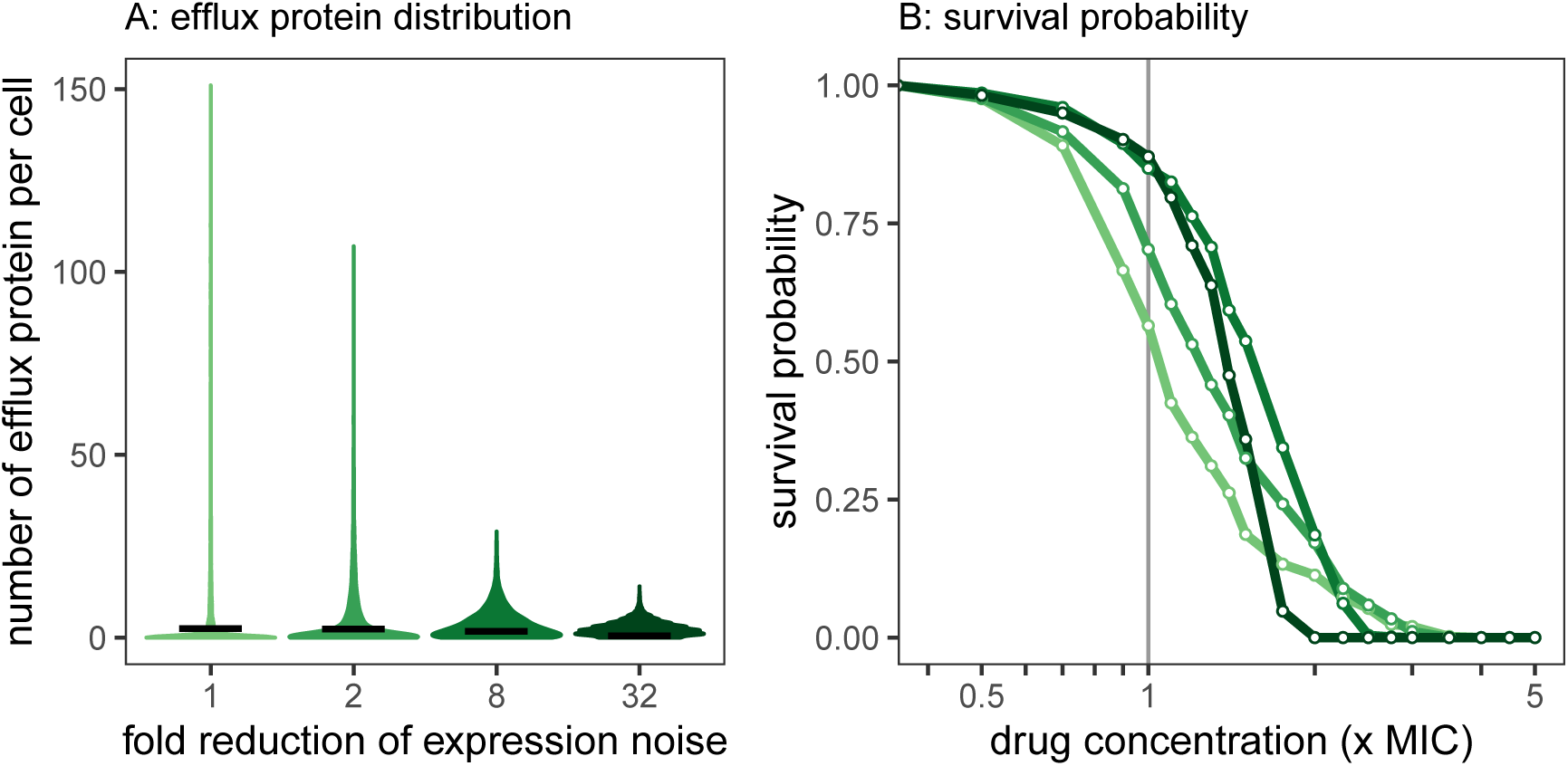
Controlling gene expression stochasticity in STRUCT-BIND mutants. A) Efflux protein number distributions across 10,000 cells with increased gene activation rate Γ_On_ and decreased translation rate *γb*. Solid black bars are the mean number of efflux protein per cell. B) Survival probability of STRUCT-BIND mutants with effect *µ* = 50, with varying expression noise (*α* ∈ {1, 2, 8, 32}, see main text). Colours correspond to the regulatory noise in panel A. Simulations are performed as in Fig. 3.

By reducing the gene expression noise (larger *α*), we note an increased IC50 but decreased IC90 (Fig. 6B), as the survival profile now resembles that of REG-ON mutants (see Fig. 3A). With reduced expression noise the efflux protein number in cells becomes more uniform and cells with extreme protein abundances disappear, making resistance homogeneous across the cell populations. Overall this shows that while noise control reduces the amount of drugs needed to inhibit the cells, the antibacterial effect also diminishes when drug concentrations fall below the MIC.

Another source of heterogeneity stems from the partitioning of proteins at cell division. Our model assumes efflux proteins are binomially distributed between two daughter cells, and so far we have only considered the symmetrical case with binomial parameter *p* = 0.5. It has been previously shown, however, that biased partitioning for efflux pumps exists and could cause long-lasting phenotypic heterogeneity (Bergmiller et al., 2017). To check the effects of partitioning bias, we varied the probability parameter *p* of the binomial distribution to favour one daughter cell with more efflux proteins at division (Supplementary Fig. S8). Among WT lineages, IC90 is unaffected since most cells have no efflux proteins, while a few cells have some with low-efficacy. In REG-ON mutants, however, we observe a significant increase in the lineage survival probability (IC90), as one cell can now accumulate a significant number of efflux proteins. In the STRUCTBIND mutants, one cell in the lineage will accumulate highly-effective efflux proteins, making it into a super-resistant cell and hence increasing the IC90. These results initially carry across to zMIC as determined from growth rate measurements – increasing the efflux distribution bias results in a higher zMIC. However, in the most extreme case (*p* = 1) where only one daughter receives all the efflux protein, the STRUCT-BIND mutant shows a decreased zMIC measurement. Here, at drug concentrations above the WT-MIC, only one cell maintains the resistance from generation to generation, and hence the maximum growth rate is zero. Our results therefore suggest that there is an optimum strategy to invest more in the fitness of one daughter which maximises the resistance of the population. This is not seen in the REG-ON mutant as almost all cells are producing efflux mRNA and proteins frequently. REG-ON daughter cells that inherited no efflux protein could replenish their resistance rapidly and survive the antibiotic, thereby contributing to population growth.

Although it is commonly accepted that increased mean expression of resistance enzymes can enhance resistance (Zwart et al., 2018), our results suggest that expression noise is also important for the population-level dynamics.

### 3.4 Survival after antibiotic pulse

The survival and extinction time properties of the different mutants are expected to affect the outcome of pulsed antibiotic treatment, which reflects the situation in a patient where drugs are not administered in a manner that maintains a constant concentration. Therefore we simulated pulsed treatments where cells are exposed to a one-time dose of antibiotic for a limited time period. For simplicity, the drug dose is expressed as a step-function without more-detailed pharmacokinetics. We then measured the survival probability for each mutant class across different treatment durations and concentrations (Fig. 7). This survival probability can also be interpreted as the genetic diversity, as it measures the fraction of unique lineages which survive the pulsed treatment.

**Figure 7:**
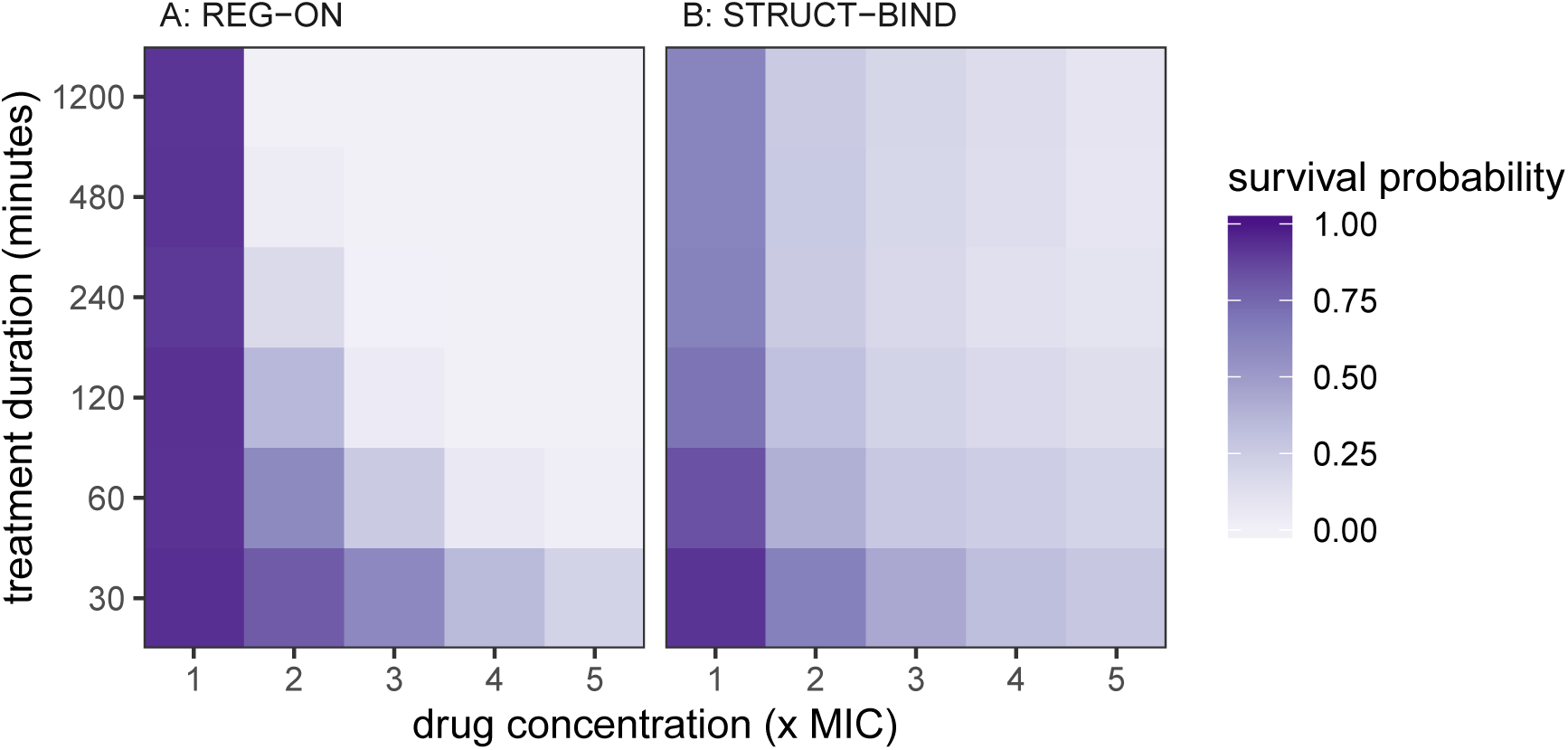
Survival probability after exposure to a pulse of antibiotics. The survival probability is indicated by colour scale. The drug dose is applied as a step-function with constant external drug concentration during the pulse, and *c*_out_ = 0 for the remainder of the experiment up to *t* = 1, 200 minutes. Simulations are performed as in Fig. 3. The mutant effect is *µ* = 200. Further results for the other cell types can be found in Supplementary Fig. S9.

REG-ON mutants have the highest survival probability at low drug concentrations (*c*_out_ ≤ 2×MIC) or with brief drug pulses (e.g. *t* = 30 minutes), as shown in Fig. 7A. This is due to their long extinction times that allow regulatory mutants to outlast drug concentrations beyond their IC90. STRUCT-BIND cells survive at higher concentrations with long drug pulses (Fig. 7B), consistent with their high IC90 values. REG-OFF mutants perform similarly to REG-ON mutants, while STRUCT-CAT, KO, and WT cells show low levels of survival (Supplementary Fig. S9).

### 3.5 Systematic analysis

So far, we find that STRUCT-BIND mutations provide most resistance, followed by REG-ON. STRUCT-CAT, however, do not increase the resistance against ciprofloxacin. For beta-lactams, it has been shown in experiments that increasing *k*_cat_ provides a high level of resistance (Knies et al., 2017; Palzkill, 2018). To check whether our results hold true for other conditions, we performed a systematic grid sampling of the model parameter space. Specifically, we vary six model parameters: binding rate, catalysis rate, diffusion rate, number of targets, number of efflux proteins, and number of drug molecules per cell at MIC, constructing 729 unique parameter combinations. For each of these we consider the WT cell, as well as the REG-ON, STRUCTBIND, and STRUCT-CAT mutants with an effect of *µ* = 200. The performance of each mutant class is quantified as their ability to increase the IC90 of the cell population relative to the WT cell. Full details of this procedure can be found in the Supplementary Material.

Through this analysis, we find that the largest effect mutations are STRUCT-BIND, but on average REG-ON outperform the other mutants (Fig. 8). REG-ON and STRUCT-BIND mutants frequently provide increases of > 10-fold in IC90, while STRUCT-BIND mutants could even reach above 100-fold in some circumstances. STRUCT-CAT mutants have little-to-no effect most of the time, but it is possible for them to have up to 10-fold advantage over the WT. Concretely, STRUCT-CAT mutants perform better when: i) the binding rate is high; ii) the average number of efflux pumps per cell is high; iii) the number of intracellular drug molecules are high; iv) the WT catalysis rate is low (Supplementary Fig. S10). In all four scenarios, catalysis becomes the rate-limiting step in the efflux of drug from the cell. In summary, while STRUCTCAT mutations could enhance resistance, their contributions are fairly limited compared to STRUCT-BIND and REG-ON.

**Figure 8:**
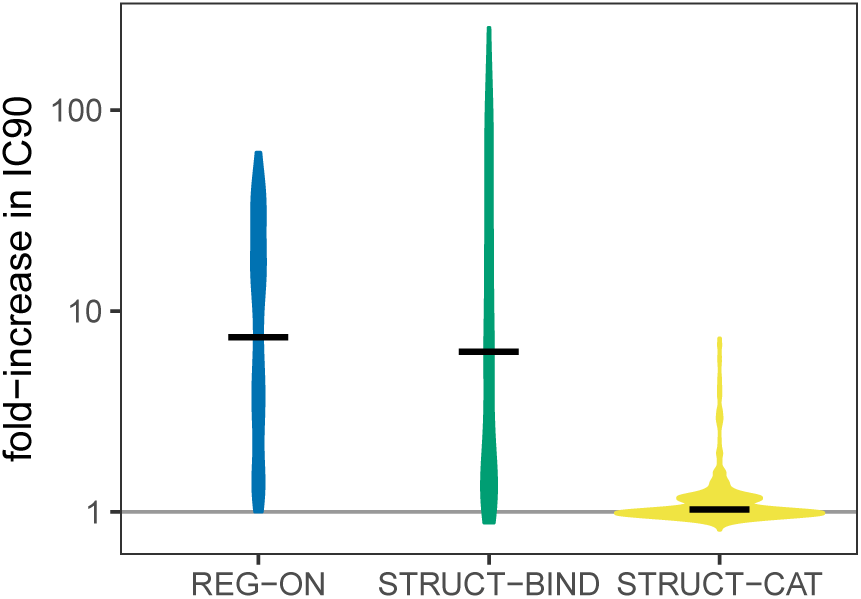
Distribution of mutant efficacy following systematic parameter sampling. We plot values of IC90 (relative to WT) across the 729 parameter combinations for REG-ON, STRUCTBIND, and STRUCT-CAT mutants with *µ* = 200. Black lines are the mean fold-increase in IC90 for each mutant across all parameter combinations. IC90 is determined from the lineage survival probability of 1,000 founder cells, as described in Fig. 3.

## 4 Discussion

Efflux pumps and drug-inactivating enzymes cause some of the most important clinical resistances (Li et al., 2015; World Health Organization, 2017). Overexpression of efflux pumps is also associated with virulence and enhanced mutation rates (Wang-Kan et al., 2017; El Meouche and Dunlop, 2018). Although stochasticity is inherent to gene expression and underlies vital cell functions, its effects on antibiotic resistance have not been systematically addressed.

To study this, we constructed an efficient agent-based model that bridges the scales between within-cell molecular processes and population-level dynamics. We considered different classes of regulatory and structural mutations that enhance resistance, and evaluated their performances under different drug pressures. Previous models have utilised drug–target binding dynamics to help explain population-level phenomena such as persistence (Abel Zur Wiesch et al., 2015). Our model extends this approach by adding enzymatic reactions. Furthermore, unlike previous deterministic models, our stochastic approach explicitly captures the noise at each step of the relevant biological processes, allowing us to dissect the effects of each source of noise on bacterial behaviour.

Regulatory mutations that increase either transcription frequency (REG-ON) or duration (REG-OFF) by the same factor have the same mean expression. Thus, a mean-field deterministic model would predict equal levels of resistance. However, increasing the duration of transcription does not reduce the noise of gene activation, leading to a subpopulation of cells with little or no resistance enzymes which remain drug-susceptible. This is not observed in the mutants with frequent gene activation, which instead show a homogeneous population of cells with equal resistance.

Structural mutations improve the biochemical properties of the resistance enzyme by increasing the binding rate to the drug (STRUCT-BIND) or the catalysis rate (STRUCT-CAT). While structural mutations may influence both binding and catalysis simultaneously, here we disentangle them to elucidate their relative importance in resistance. We find that improving the binding rate has a significant effect on resistance, while improving the catalysis rate may have no effect (at least for our parametrisation of *E. coli* exposed to ciprofloxacin).

To understand the robustness of these conclusions, we systematically investigated a large range of parameter space taking into account different target, efflux, and drug dynamics. Increasing resistance through faster catalysis (STRUCT-CAT) is effective when the number of efflux enzymes is greater than or equal to the number of targets, which may be the case when considering beta-lactamase resistance where increased catalytic rates have been detected (Knies et al., 2017; Palzkill, 2018) in mutants with large increases in MIC (Hall, 2002; Schenk et al., 2012). Furthermore, STRUCT-CAT mutants perform better when the binding rate is high or when the WT catalysis rate is low, such that catalysis is rate limiting. Our findings, however, suggest that drug binding is predominantly the rate-limiting step in the efflux of antibiotics from the cell.

In general, high regulatory noise results in heterogeneity in phenotypic resistance whereas low noise results in homogeneous resistance. By controlling the gene-expression noise, we show that noise reduction may facilitate bacterial inhibition of mutants by reducing their IC90. At the same time, mutants also have increased IC50, suggesting that drugs could be even less efficient when falling into sub-MIC concentrations. This is relevant for exploring the clinical potential of treatments which modulate gene-expression noise. Such noise-modulating chemicals have, for example, recently shown promising effects on reactivating HIV from latency, a process that relies on high gene expression noise (Dar et al., 2014). The success of HIV-latency modulators has provided a new concept in drug discovery and we envision that a similar approach may be tested to regulate resistance in bacteria.

In clinical antibiotic treatments the drug concentration is not maintained at a consistently high level, as opposed to laboratory experiments. Here we assume a constant drug scenario as a minimalist approach for two reasons: i) computational efficiency as cells can be independently simulated; and ii) if the cells can modify the drug concentration, then we would also have to account for spatial variation of this concentration, which in turn would require biomechanical models of cell division and movement. The implementation of such a complex model would cloud the inferences we make from cellular stochasticity. On the other hand, the constant-concentration assumption is only valid when molecule numbers are high and when local fluctuations in the drug concentration are negligible. The latter may not apply if pharmacokinetics exist or if drugs are deactivated by the bacteria. One specific scenario that makes our assumption valid is if we assume the media is well-mixed: i.e. the diffusion speed of the drug in the media is much faster than the diffusion across the cell membranes. If the volume of media is much greater than the total volume of all cells in the media, then the external drug concentration is approximately constant across the experiment. Further complications would be the release of drug upon cell lysis, which again would not be a problem in the aforementioned large-volume, high-density scenario.

While working with constant drug concentrations, we varied the duration of the drug dose to study how pulsed treatments affect bacterial survival. This is relevant as the mutant classes show different extinction times (Supplementary Fig. S6), which could affect treatment success. Regulatory mutants have long survival times even at high concentrations, which leads to high survival probabilities as long as the drug is applied in a brief pulse. Thus, regulatory mutants could be more clinically problematic than structural mutants despite having lower IC90. This outcome will not be reflected in standard experimental protocols based on constant concentrations such as MIC-measurements. Therefore, a comprehensive understanding of mutant dynamics requires more detailed assessment methods for cell death and growth across different antibiotic concentrations.

Our drug–target model is based on ciprofloxacin, gyrase, and the AcrAB-TolC efflux pump. This system has the advantage of relatively few drug targets per cell and drug that acts at low concentrations, which reduces computation time. We parametrised our model based on previous experimental findings. It can accurately reproduce cell population dynamics under various drug concentrations as well as within-cell protein distributions under stochastic expression (Fig. 4E, Supplementary Fig. S4B). We therefore expect our model and its findings to be qualitatively robust.

Given that stochastic gene expression is a general mechanism in cell biology, our findings may also offer insights into general bacterial adaptation to changing environments. One example is the evolutionary reproducibility of adaptation via gain-of-function mutations: in certain biological systems, regulatory mutations seemingly take precedence over structural mutations and vice versa (Toprak et al., 2012; Blank et al., 2014; Lind et al., 2015). Here we show that stochastic gene expression has temporal and heterogeneous effects on regulatory versus structural mutants, which helps explain why certain mutations could be advantageous.

In conclusion, the dynamics of gene expression could have a strong impact on the efficacy, and therefore the evolution, of resistance mutations. In particular, the intrinsic stochasticity of gene expression is a crucial determinant of evolutionary success. The framework we developed in this study now opens further opportunities to assess the impact of stochastic gene expression on bacterial adaptation, and the impact of molecular biology on evolution by extension.

## 5 Supplementary material

Supplementary Tables S1-S3 and Supplementary Figs. S1-S10 can be found in the Supplementary Material. Also included is the description of the mean-field model, the comparison of different MIC measurements, and the description of the systematic parameter search. All code and generated data can be found at the github repository https://github.com/ashcroftp/StochasticGeneExpression2019. This will be archived on Zenodo and made publicly available prior to publication.

## 6 Acknowledgments

We thank Pia Abel zur Wiesch for commenting on an earlier version of the manuscript. This work was supported by SystemsX.ch, MRD Project 2014/266 412 “StemSysMed” (PA & SB), grant no. 31003A 169978 from the Swiss National Science Foundation (MA), grant no. 310030B 176401/1 from the Swiss National Science Foundation (SB), and grant no. 268540 from the European Research Council (SB).

## Supplementary Material

### S.I Mean field model

From the reactions in Table 2 of the manuscript, we can construct a mean-field ordinary differential equation (ODE) model:

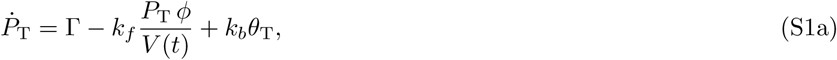

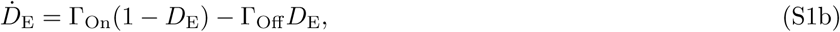

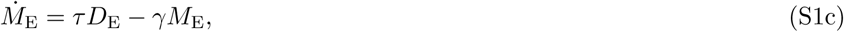

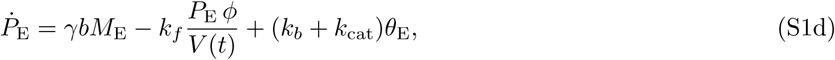

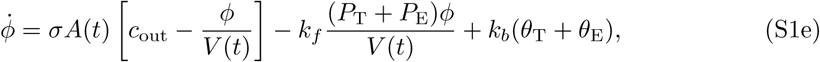

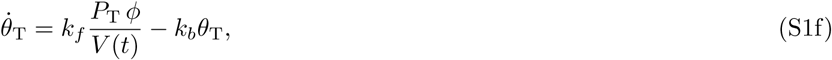

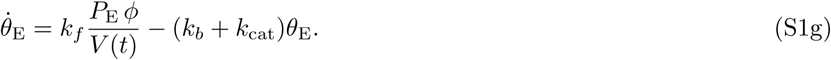

For the behaviour shown in Fig. 2 of the manuscript, these equations are integrated until time *t*_*G*_, after which the molecule numbers are halved to represent dilution due to cell division.

### S.II Comparing MIC measurements

The population of cells roughly follows the growth law *n*(*t*) = *n*(0)10^*Ψt*^, where *n*(0) is the initial population size and *Ψ* is the log-10 growth rate. To find a population growth rate, we then solve this equation for *Ψ*, i.e.,

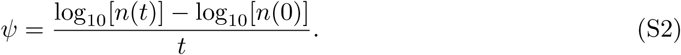

If we measure a lineage extinction probability of *X* (say, 90%), then the maximum population size at time *t* that we could observe is 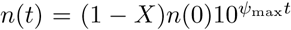, where each of the surviving lineages (1 − *X*) grow at the maximum rate. Likewise, the minimum population size with a lineage extinction probability of *X* is *n*(*t*) = (1 − *X*)*n*(0), i.e. each surviving lineage only has a single cell remaining. We therefore have the following bounds for the growth rate:

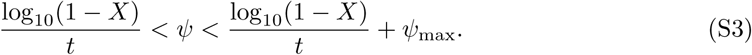

These bounds are highlighted in Supplementary Fig. S1A, from which we see that IC90 measurements after one hour would overestimate the zMIC – instead the IC90 would correspond to a negative net growth rate of the population. When measuring growth rates at later timepoints, we can have correspondence between the IC90 and zMIC, while IC50 is likely to underestimate the concentration which gives a net growth rate of zero.

To confirm the above findings, we compare the ICX values extracted from lineage survival simulations with inhibitory concentrations as measured in growth rate simulations. From the overnight lineage survival probability simulations, we interpolate IC50 (50% lineage survival), IC90 (10% lineage survival), and IC99 (1% lineage survival). From the five hour growth rate simulations, we interpolate the drug concentrations at which *Ψ* = 0.1*Ψ*_max_ (10% max growth rate), *Ψ* = 0.01*Ψ*_max_ (1% max growth rate), and *Ψ* = 0 (zero growth, or zMIC). We measure the growth rates at multiple timepoints to determine the observations are dependent on experiment duration. In Supplementary Fig. S1B, we see that the inhibitory concentrations inferred from growth rates converge to IC90 for the REG-ON, REG-OFF, STRUCT-CAT, WT and KO cell types, and this convergence occurs within 3 hours. IC50, on the other hand, always underestimates the true inhibitory concentration. For the STRUCT-BIND and REG-BURST mutants, the inhibitory concentration interpolated from growth rates continually increases with time and IC90 underestimates the zMIC. Therefore IC90 is a conservative estimate of the MIC of the STRUCT-BIND and REG-BURST mutants. In summary, we find that IC90 is the most suitable measure of MIC which generally correlates well with the zMIC drug concentration.

### S.III Systematic simulations

Results in the manuscript are based on a single resistance mechanism, the AcrAB-TolC efflux pump, providing resistance to a specific drug, ciprofloxacin. To shed more light on our results, we conduct a systematic sweep of the parameter space to assess in which regimes different mutant classes are advantageous, and what are the maximum levels of resistance that can be observed.

We consider all permutations of the parameters listed in the table below:

**Table S1:** Summary of parameter values. ^a^ Average values per generation. ^b^ Number of efflux pumps is increased by increasing the mRNA burst size *b* (DNA activation/deactivation and mRNA degradation rates kept constant). ^c^ Evaluated at 1×MIC.

| Parameter | Values |
| --- | --- |
| Binding rate ($\text{M}^{-1}\text{s}^{-1}$) | $10^4, 10^5, 10^6$ |
| Catalysis rate ($\text{s}^{-1}$) | $10^{-4}, 10^{-2}, 10$ |
| Drug diffusion rate ($\text{ms}^{-1}$) | $10^{-11}, 10^{-10}, 10^{-9}$ |
| Number of targets<sup>a</sup> | $10, 10^2, 10^3$ |
| Number of efflux pumps<sup>a,b</sup> | $10, 10^2, 10^3$ |
| Number of drug molecules<sup>a,c</sup> | $10, 10^2, 10^3$ |

The three values for each of the six parameters gives a total of 3^6^ = 729 unique parameter combinations. For each of these we consider the WT cell, as well as the REG-ON, STRUCTBIND, and STRUCT-CAT mutants with an effect of *µ* = 200.

For each parameter combination, we need to determine the external drug concentration for which the average number of drug molecules per WT cell corresponds to the value in the table above. This concentration is then the MIC of that specific parameter combination. We then use the mean-field model [Eq. (S1)] to calculate the average fraction of bound targets per cell, and we assign this value as the MIC fraction, *ρ*_MIC_. We set *κ* = 3 throughout, and the remaining parameters are the same as in Supplementary Table S2.

Concretely, for each parameter combination we do the following:

1. Load the basic model parameters, and modify the binding, catalysis, and diffusion rates, and the number of targets directly;
2. Set the rate of efflux mRNA transcription such that the average number per cell matches the designated value;
3. Using these parameters, we integrate the mean-field equations for different external drug concentrations, and solve for an external concentration which gives the average number of drug molecules per cell as given in the table above;
4. We integrate the mean-field equations at this given concentration, and determine the MIC fraction as the average fraction of bound targets.

With this procedure, each of the 729 combinations can be rapidly parametrised, although not completely accurately. We then compute the IC90 values for the each of the 729×{WT, REGON, STRUCT-BIND, STRUCT-CAT} combinations, and extract the relative increase in IC90 of each of the mutant cells. The distribution of these mutational effects are shown in Fig. 8 of the manuscript.

To understand when each mutant is advantageous, we can compute the correlation between each of the six parameters and the fold-increase in IC90. This is shown in Fig. S10. A general trend across the three mutants is that resistance is highest when the baseline number of efflux proteins is high, binding rates are fast and diffusion rates are slow.

### S.IV Supplementary tables

Model parameters are given in Table S2, and drug-specific parameters are given in Table S3. IC50 and IC90 values for ciprofloxacin treatments can be found in Table S4.

**Table S2:**
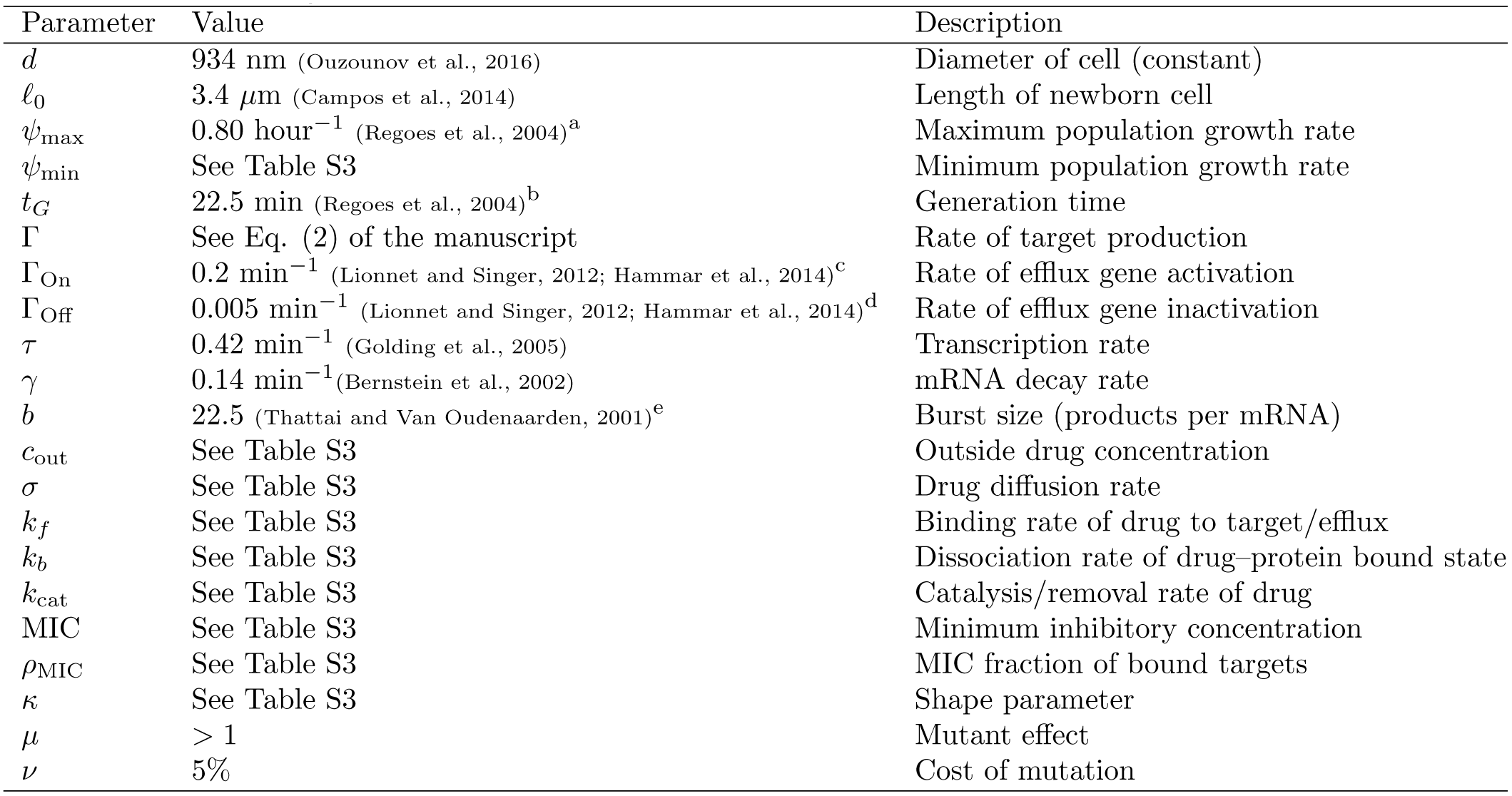
Summary of parameter values. ^a^Weighted average of reported values. ^b^*t*_*G*_ = log(2)*/*(log(10)*Ψ*_max_). ^c^Inactive time reported as 0.5–3,000 min. ^d^Active time reported as 5–60 min. ^e^Examples range from 5–40.

**Table S3:** Drug-specific parameters extracted from the literature. ^a^Value is based on AcrB efflux pump. ^b^Values chosen according to Fig. S2. ^c^Values chosen according to Fig. S3.

| Drug | Ciprofloxacin | Rifampicin |
| --- | --- | --- |
| Target | DNA Gyrase | RNA polymerase |
| Number of targets | 300 (Chong et al., 2014; Stracy et al., 2019) | 11,400 (Abel Zur Wiesch et al., 2015) |
| MIC ($\mu\text{g ml}^{-1}$) | 0.03 (Regoes et al., 2004) | 8.00 (Regoes et al., 2004) |
| Molar mass ($\text{g mol}^{-1}$) | 331.347 | 822.953 |
| Diffusion rate ($\text{m s}^{-1}$) | $2.0 \times 10^{-11}$ (Abel Zur Wiesch et al., 2015) | $2.0 \times 10^{-11}$ (Abel Zur Wiesch et al., 2015) |
| Binding rate ($\text{M}^{-1} \text{s}^{-1}$) | $3.6 \times 10^4$ (Kampranis and Maxwell, 1998) | $1.2 \times 10^6$ (Abel Zur Wiesch et al., 2015) |
| Dissociation rate ($\text{s}^{-1}$) | $3.0 \times 10^{-4}$ (Kampranis and Maxwell, 1998) | $1.2 \times 10^{-3}$ (Abel Zur Wiesch et al., 2015) |
| Catalysis rate ($\text{s}^{-1}$) | 10.0 (Nagano and Nikaido, 2009)<sup>a</sup> | 10.0 (Nagano and Nikaido, 2009)<sup>a</sup> |
| Min. pop. growth rate ($\text{h}^{-1}$) | -6.5 (Regoes et al., 2004) | -4.3 (Regoes et al., 2004) |
| Shape ($\kappa$) | 2.0<sup>b</sup> | 2.0<sup>c</sup> |
| $\rho_{\text{MIC}}$ | 0.081<sup>b</sup> | 0.294<sup>c</sup> |

**Table S4:**
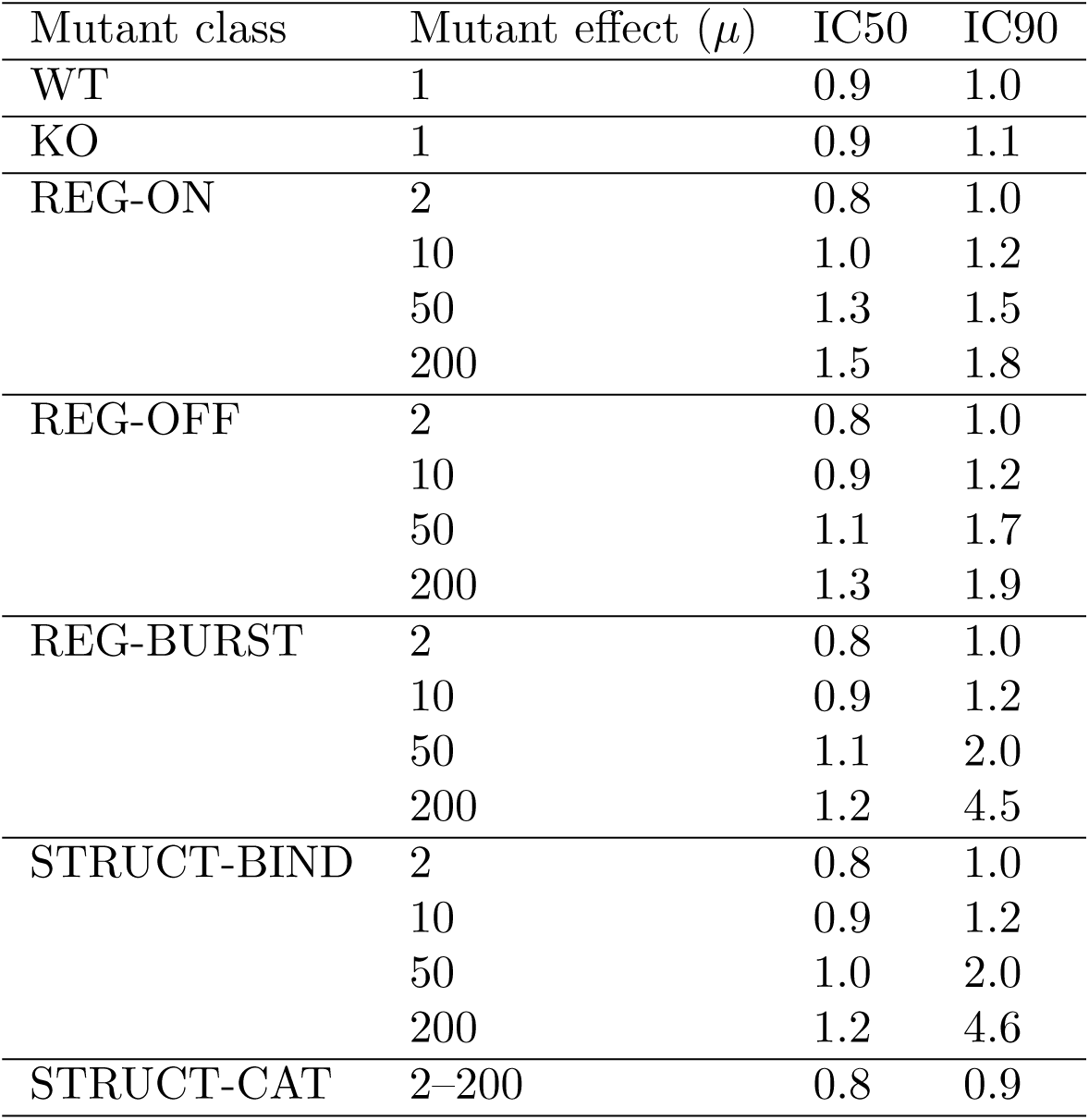
Profiles of resistance to ciprofloxacin. The IC50 and IC90 values are expressed as multiples of the MIC for the WT strain. These values are extracted from Fig. 3 of the manuscript using spline interpolation, and are defined as the drug concentrations at which 50% and 90% of the lineages from the initial bacterial inoculum are killed.

| Mutant class | Mutant effect ($\mu$) | IC50 | IC90 |
| --- | --- | --- | --- |
| WT | 1 | 0.9 | 1.0 |
| KO | 1 | 0.9 | 1.1 |
| REG-ON | 2 | 0.8 | 1.0 |
|  | 10 | 1.0 | 1.2 |
|  | 50 | 1.3 | 1.5 |
|  | 200 | 1.5 | 1.8 |
| REG-OFF | 2 | 0.8 | 1.0 |
|  | 10 | 0.9 | 1.2 |
|  | 50 | 1.1 | 1.7 |
|  | 200 | 1.3 | 1.9 |
| REG-BURST | 2 | 0.8 | 1.0 |
|  | 10 | 0.9 | 1.2 |
|  | 50 | 1.1 | 2.0 |
|  | 200 | 1.2 | 4.5 |
| STRUCT-BIND | 2 | 0.8 | 1.0 |
|  | 10 | 0.9 | 1.2 |
|  | 50 | 1.0 | 2.0 |
|  | 200 | 1.2 | 4.6 |
| STRUCT-CAT | 2-200 | 0.8 | 0.9 |

### S.V Supplementary figures

Comparison of MIC measurements is shown in Supplementary Fig. S1. Parameter screening for ciprofloxacin is shown in Supplementary Fig. S2. Parameter screening for rifampicin is shown in Supplementary Fig. S3. Molecule distributions are shown in Supplementary Fig. S4. Results for REG-BURST mutants are shown in Supplementary Fig. S5. Extinction times are shown in Supplementary Fig. S6. Time-kill curves are shown in Supplementary Fig. S7. Results of biasing the binomial distribution of efflux pumps is shown in Supplementary Fig. S8. Survival probabilities following pulsed drug treatment are shown in Supplementary Fig. S9. The correlation between parameter values and mutational effects from the systematic grid sampling is shown in Supplementary Fig. S10.

**Figure S1:**
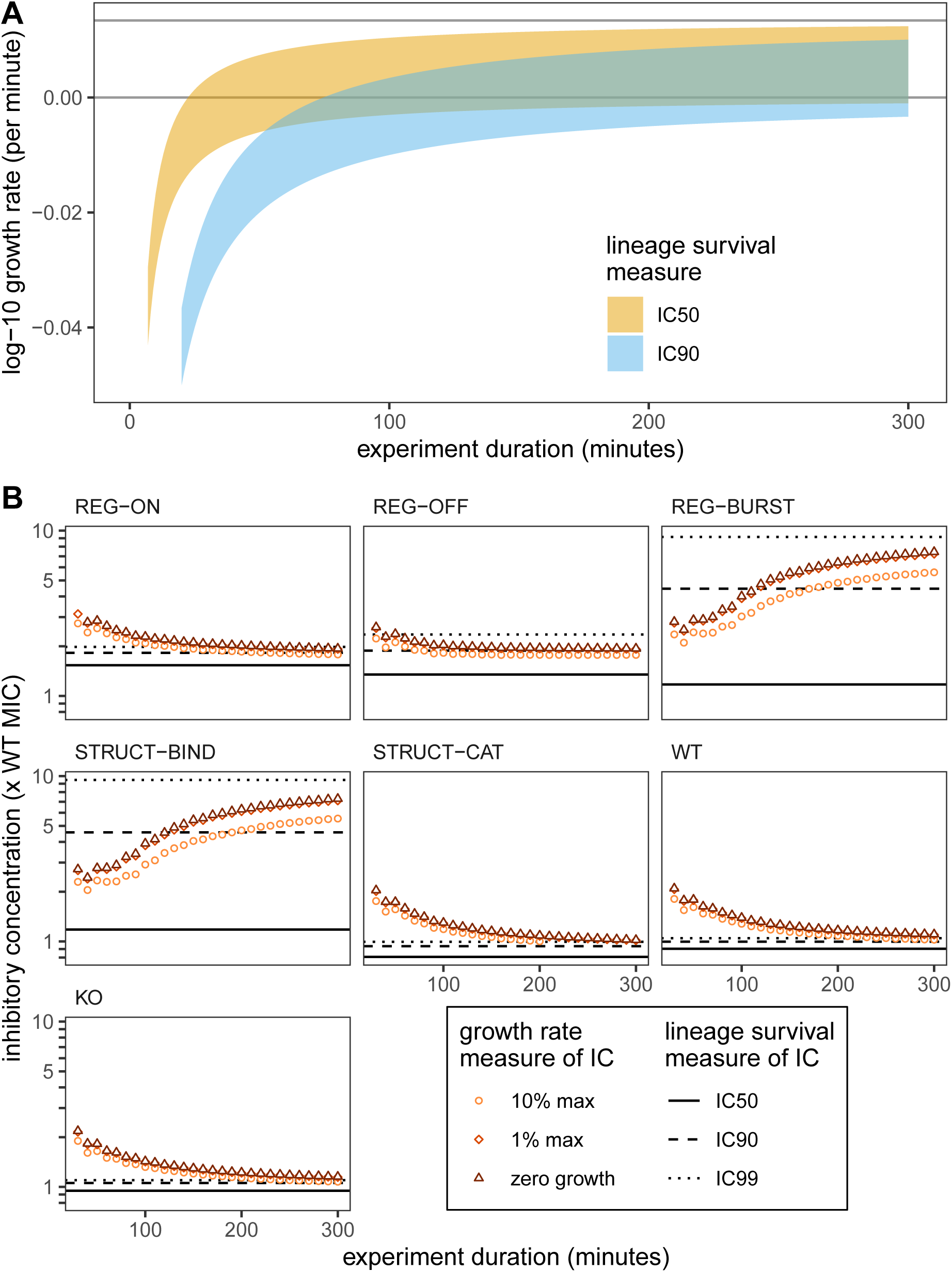
Comparing MIC measurements from growth rate data and lineage survival probabilities. A) Range of growth rates that can be measured for a given lineage survival probability. IC50 has a survival probability of 50%, while IC90 has a survival probability of 10%. Curves are predicted from Eq. (S3). Grey horizontal lines are *Ψ*_max_ (upper) and zero growth rate. B) Comparison of inhibitory concentrations as determined by lineage survival (lines) or growth rates (symbols). Here we use the parameters for ciprofloxacin. Lineage survival is measured after 1,200 minutes, while growth rates are computed as described in Fig. 4 of the manuscript.

**Figure S2:**
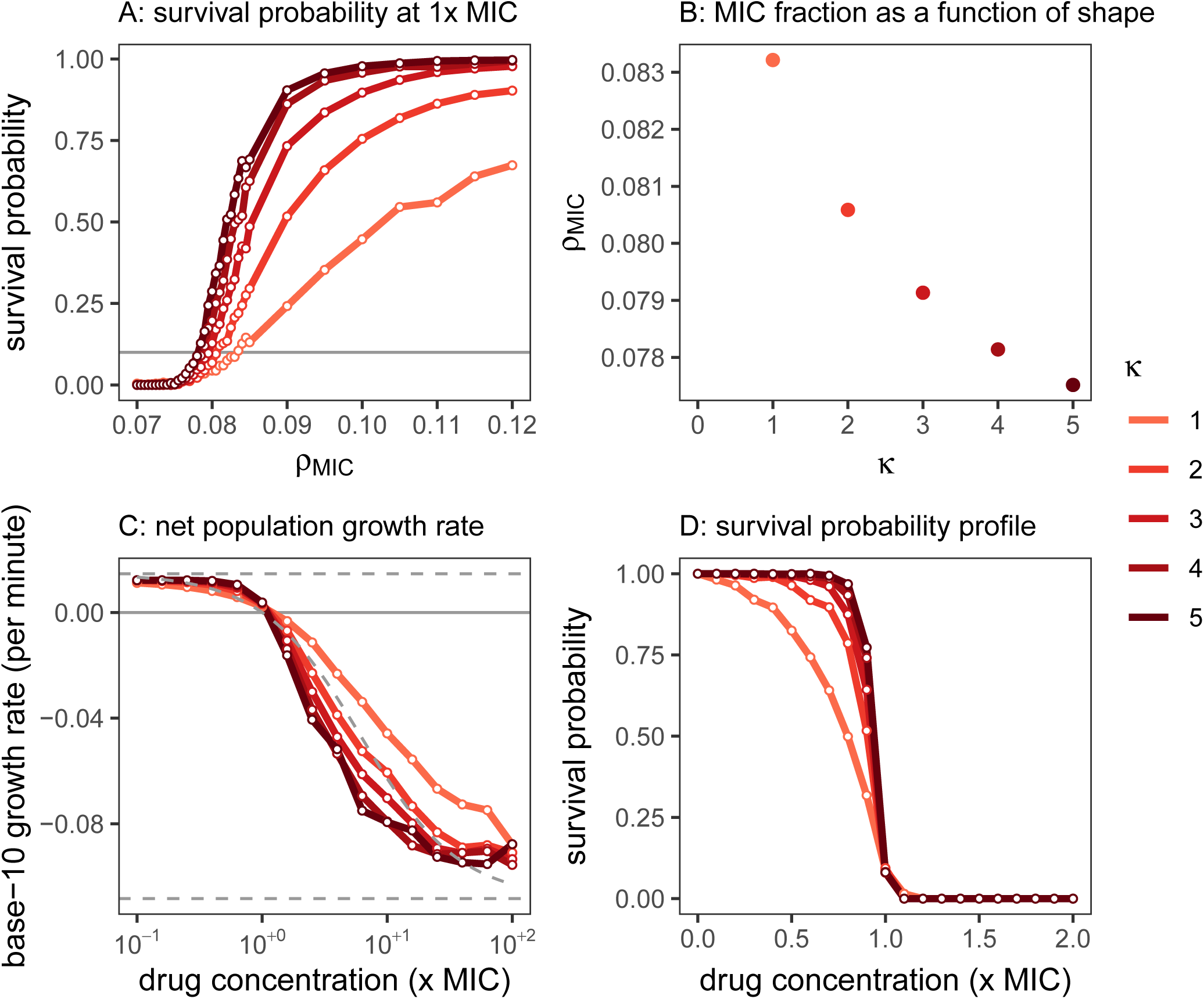
Parameter screening for ciprofloxacin. A) We calculate the survival probability for 1,000 lineages initiated from a single cell at 1×MIC while varying the MIC fraction of bound targets (*ρ*_MIC_) and the shape parameter *κ*. B) From panel A we interpolate the value of *ρ*_MIC_ which gives a survival probability of 10% (IC90). C) For each pair of *κ* and *ρ*_MIC_ in panel B, we simulate a population of 100,000 cells under different drug pressure for three hours, and extract the net population growth rate. This can be compared with the maximum and minimum reported rates (dashed horizontal lines), and the growth curve reported by Regoes et al. (2004) (dashed curve). D) For each pair of *κ* and *ρ*_MIC_ in panel B, we calculate the survival probability of 1,000 lineages initiated from a single cell under different drug pressure. From this figure we determine the shape parameter for the death rate of ciprofloxacin is *κ* = 2 and the MIC fraction is *ρ*_MIC_ = 0.081.

**Figure S3:**
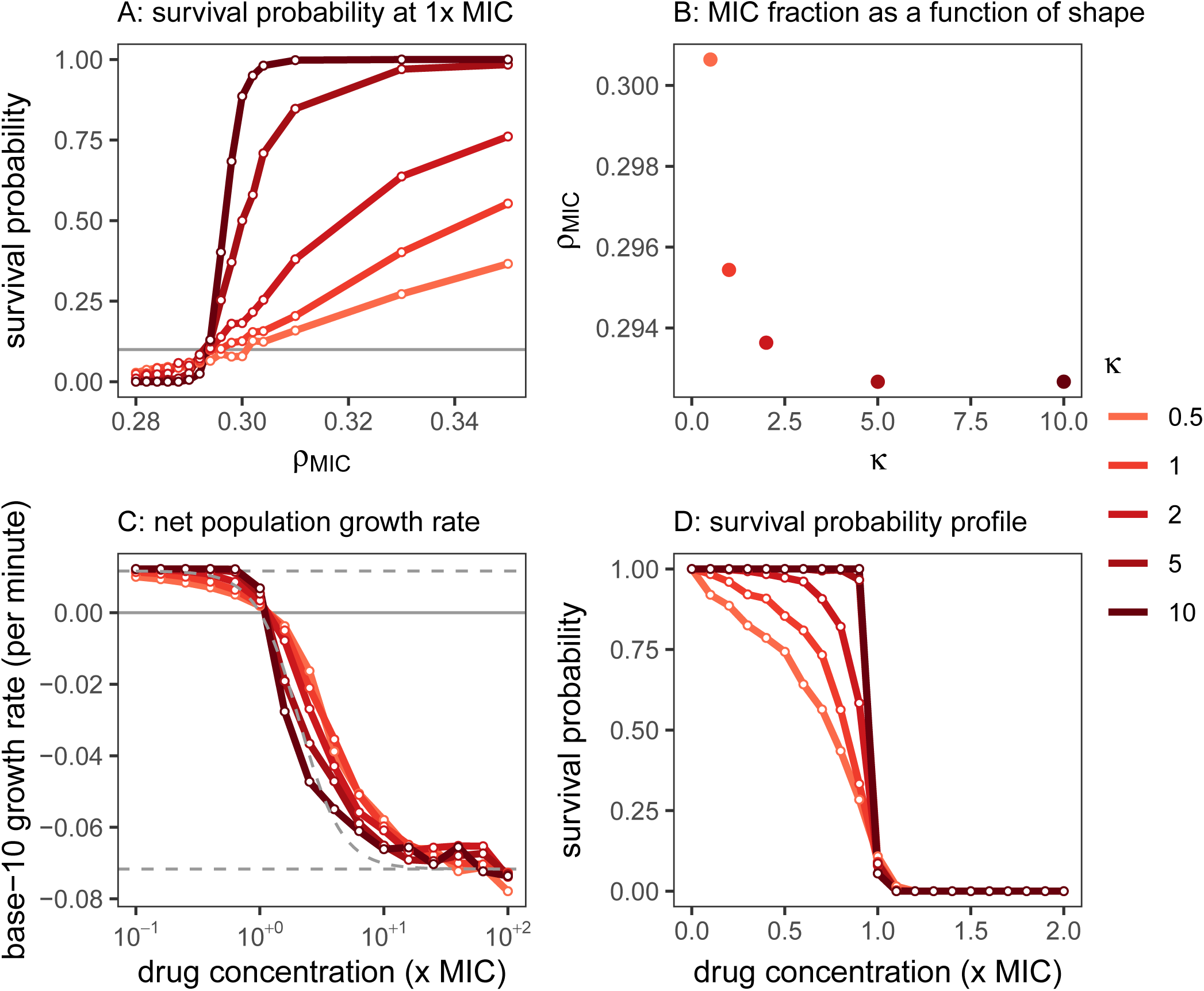
Parameter screening for rifampicin. A) We calculate the survival probability for 1,000 lineages initiated from a single cell at 1×MIC while varying the MIC fraction of bound targets (*ρ*_MIC_) and the shape parameter *κ*. B) From panel A we interpolate the value of *ρ*_MIC_ which gives a survival probability of 10% (IC90). C) For each pair of *κ* and *ρ*_MIC_ in panel B, we simulate a population of 100,000 cells under different drug pressure for three hours, and extract the net population growth rate. This can be compared with the maximum and minimum reported rates (dashed horizontal lines), and the growth curve reported by Regoes et al. (2004) (dashed curve). D) For each pair of *κ* and *ρ*_MIC_ in panel B, we calculate the survival probability of 1,000 lineages initiated from a single cell under different drug pressure. From this figure we determine the shape parameter for the death rate of rifampicin is *κ* = 2 and the MIC fraction is *ρ*_MIC_ = 0.294.

**Figure S4:**
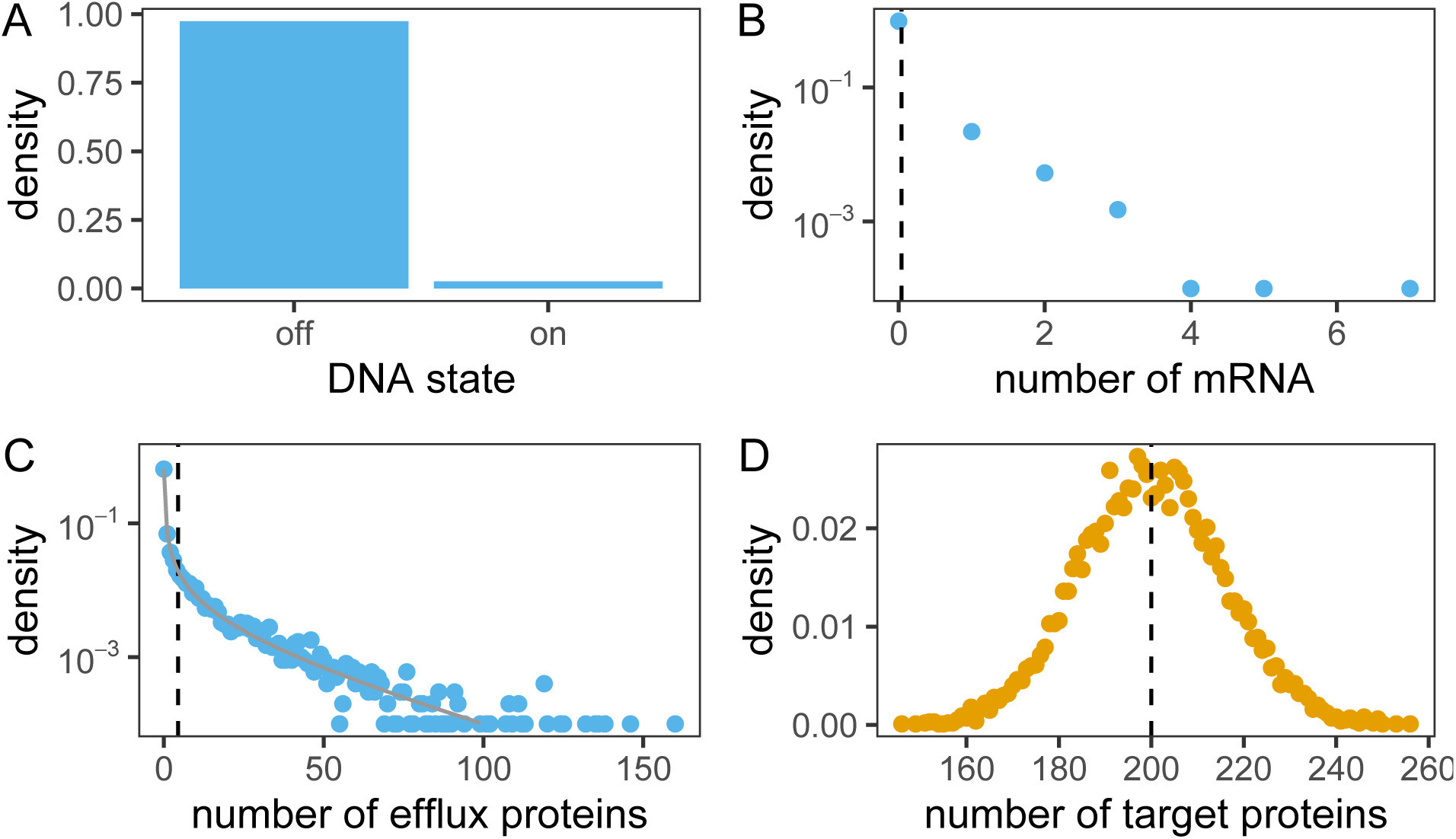
The distribution of efflux and target states across 10,000 cells immediately after cell division. These are sampled after *t* = 1, 200 minutes in the absence of drugs. The panels show: A) expression state of the efflux gene; B) distribution of efflux mRNA across the population of cells; C) number of efflux proteins per cell, which fits to a negative binomial distribution (grey line); D) number of target proteins (gyrase) per cell. Parameters are based on the targets of ciprofloxacin, and can be found in Tables S2 and S3.

**Figure S5:**
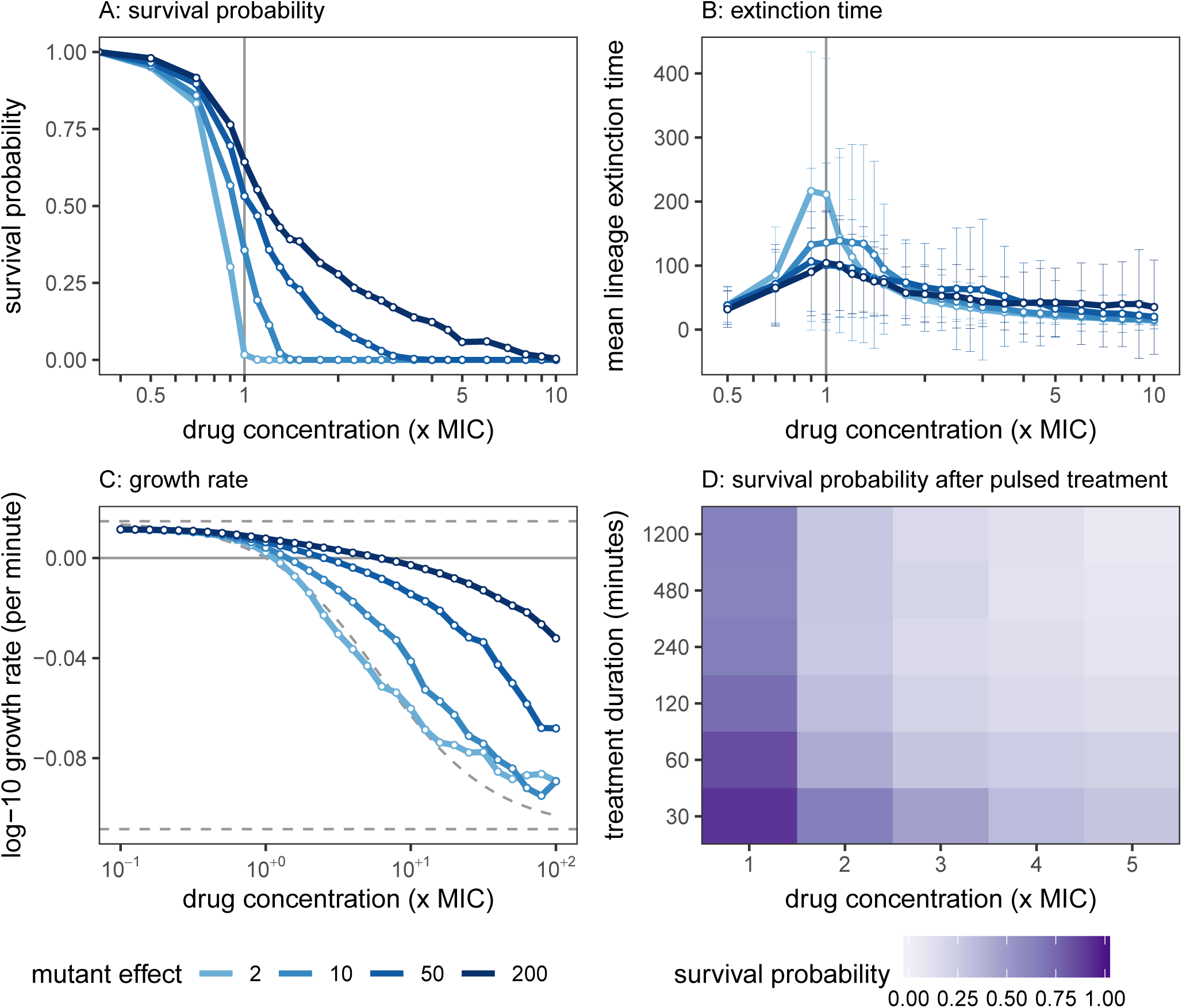
The dynamics of the REG-BURST mutants are very similar to those of STRUCTBIND. Survival probability (A) and extinction times (B) of the REG-BURST mutants across drug concentrations are calculated as in Fig. 3 of the manuscript and Fig. S6. Colour scale indicates the mutant effect (*µ*). C) The net population growth rates as computed in Fig. 4 of the manuscript. D) Survival probability after exposure to a pulse of antibiotics for the REG-BURST mutant with 200-fold effect. The survival probability is indicated by colour scale. Simulations are performed as in Fig. 7 of the manuscript and Fig. S9.

**Figure S6:**
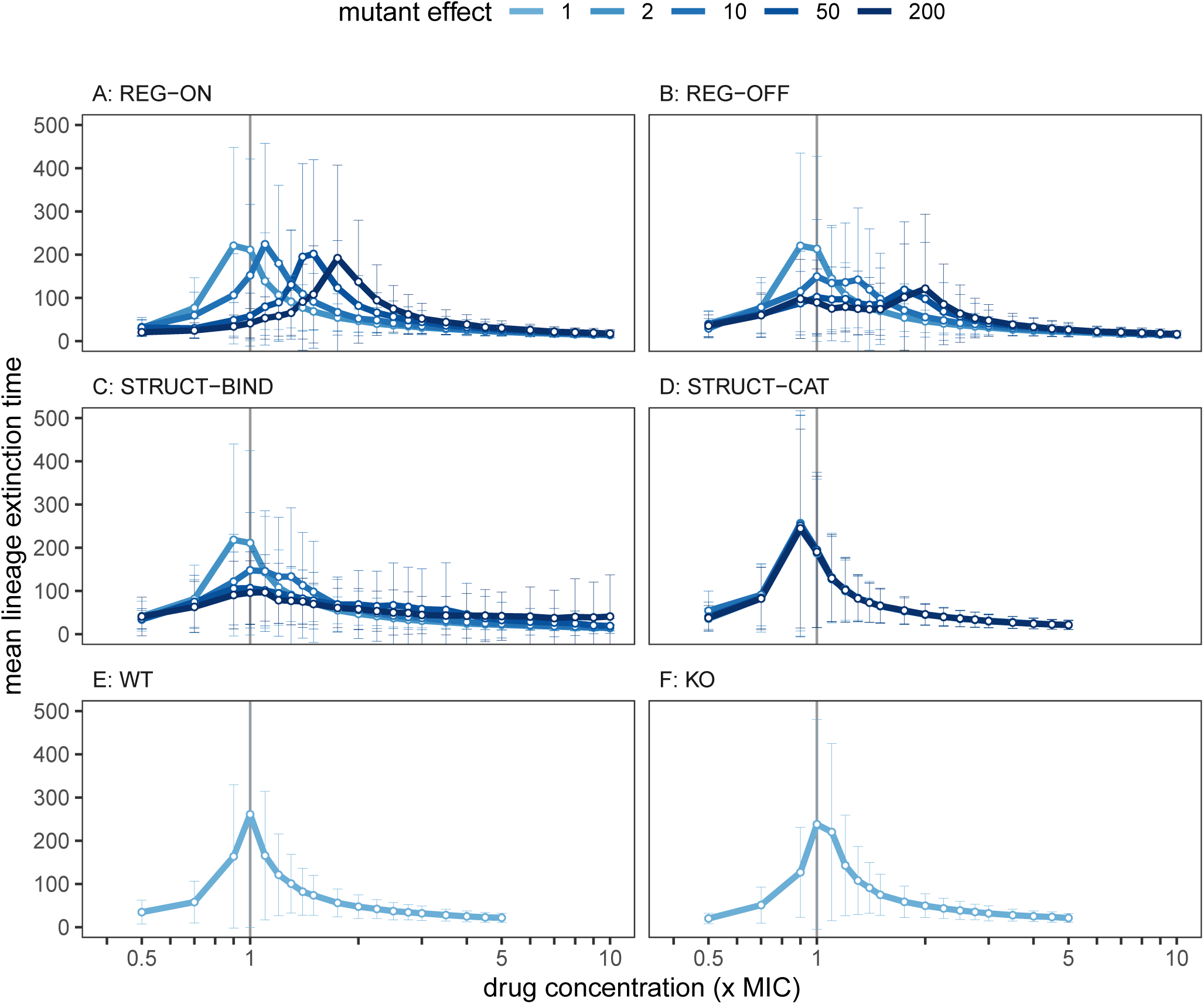
Mean extinction times of lineages of the different mutant classes. These are extracted from the same data as Fig. 3 of the manuscript. Error bars indicate the standard deviation of the extinction times.

**Figure S7:**
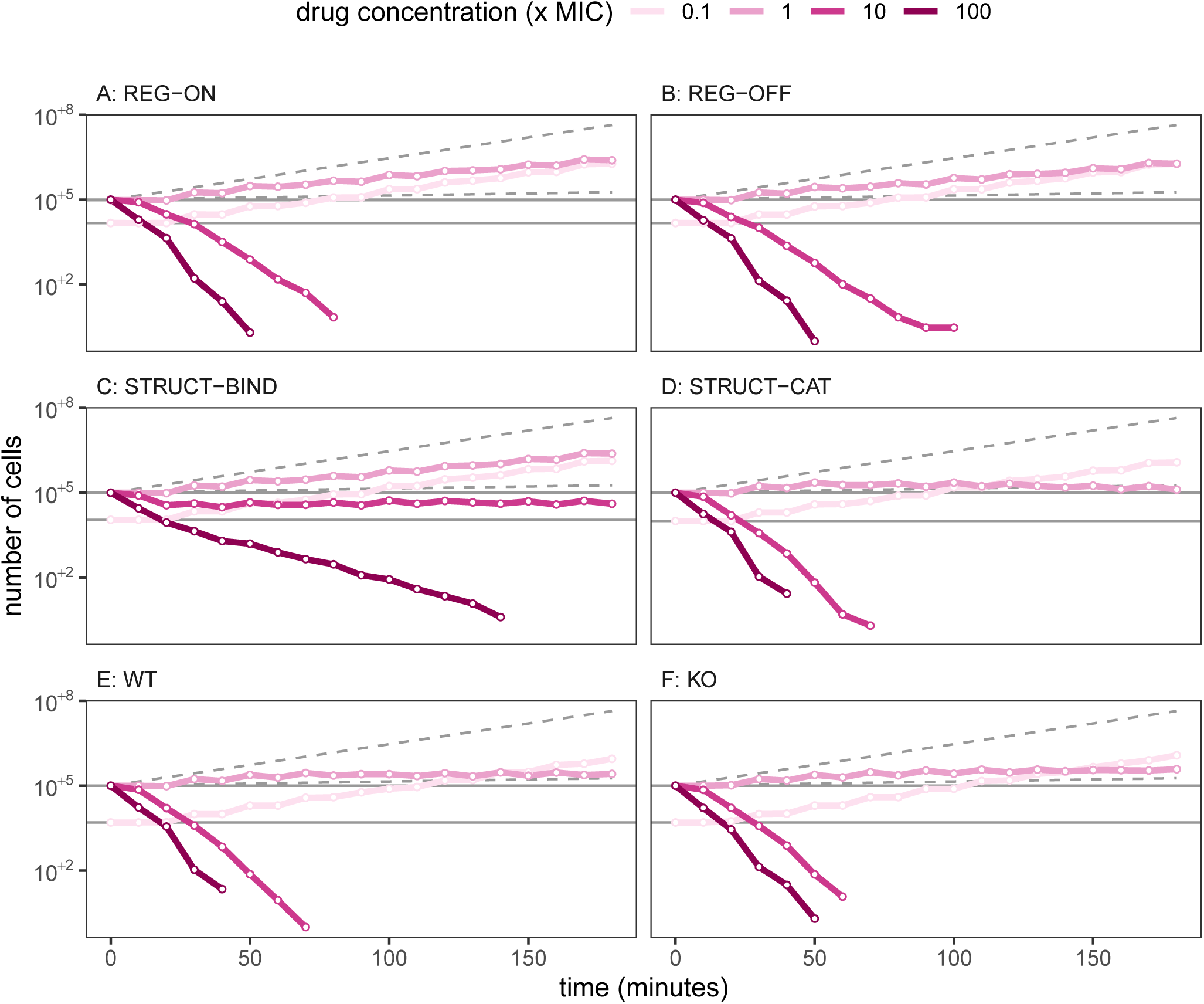
Time-kill curves, from which the growth rates in Fig. 4 of the manuscript are extracted. Here we only show mutant effect parameters *µ* = 200 (for REG-ON, REG-OFF, STRUCT-BIND, and STRUCT-CAT), and a subset of ciprofloxacin concentrations. Upper dashed line is the projected growth trajectory in the absence of drug, based on the maximum growth rate reported by Regoes et al. (2004). Lower dashed line is the trajectory if the growth rate is reduced by 90% from its maximum value. Horizontal line represents the zero growth scenario. Note that at very low drug concentrations, we reduced the initial size of the population from 10^5^ to ∼ 10^4^ due to the computational cost. This has no effect on the measurement of growth rates as the population size never approaches small numbers at these drug concentrations.

**Figure S8:**
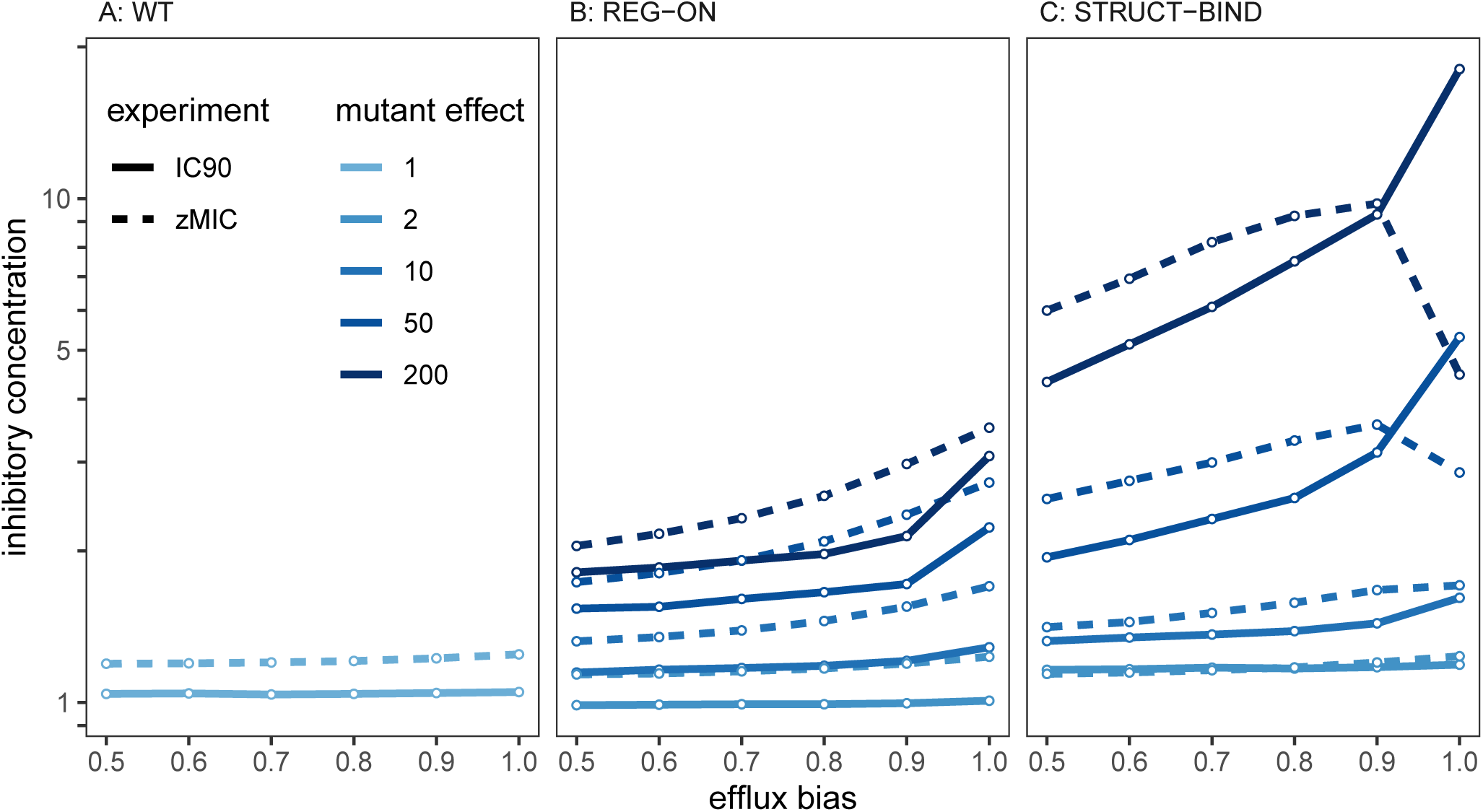
Impact of varying the efflux bias distribution during cell division. The efflux proteins (both bound and unbound) are distributed among the two daughter cells according to a binomial distribution with parameter *p*, while remaining molecules are distributed with parameter *p* = 0.5. To extract IC90 values, we compute the lineage survival probability of 1,000 initial cells over a range of drug concentrations (*c*_out_ ∈ [0, 10]×MIC), mutant effects (*µ* ∈ [1, 200]), and efflux bias parameters (*p* ∈ [0.5, 1.0]), for *t* = 1, 200 minutes. For zMIC values, we simulated up to 100,000 initial cells and tracked the population size for *t* = 180 minutes over the similar parameter ranges (*c*_out_ ∈ [10^−1^, 10^2^]) before extracting the growth rates as in Fig. 4 of the manuscript.

**Figure S9:**
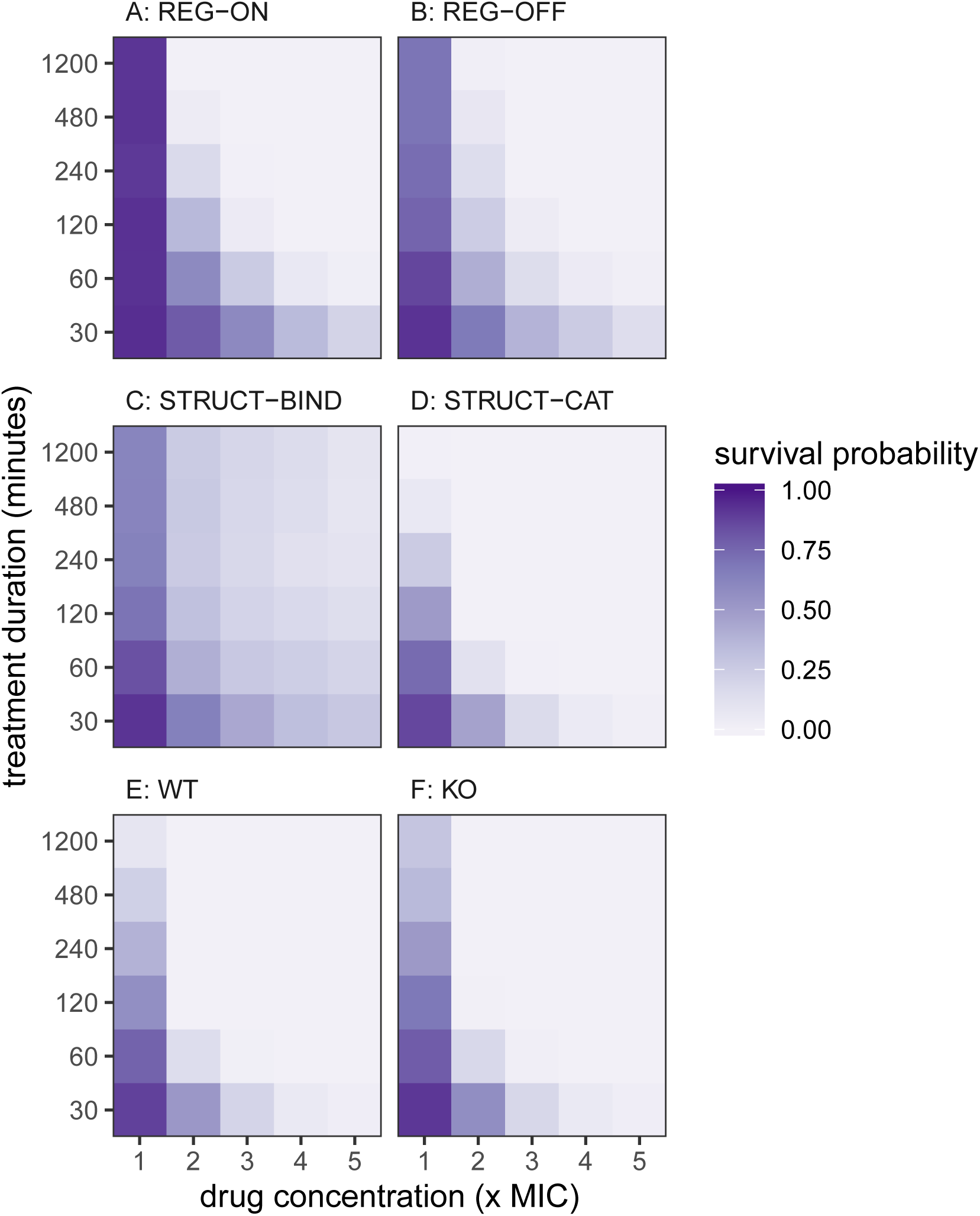
Survival probability after exposure to a pulse of antibiotics. The survival probability is indicated by colour scale. The drug dose is applied as a step-function with constant external drug concentration during the pulse, and *c*_out_ = 0 for the remainder of the experiment up to *t* = 1, 200 minutes. Simulations are performed as in Fig. 3 of the manuscript. The mutant effect is *µ* = 200.

**Figure S10:**
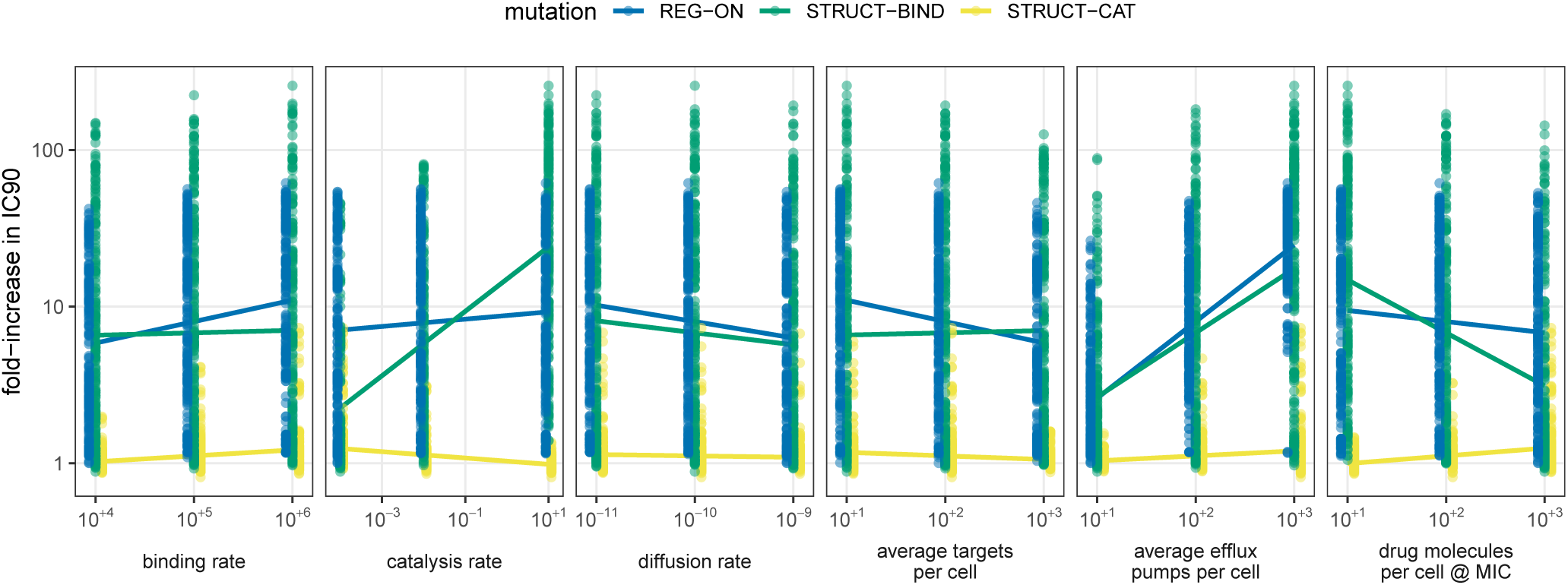
Correlation between parameter values and mutant efficacy following systematic parameter sampling. IC90 is measured for each of the 729 parameter combinations in WT, REGON, STRUCT-BIND and STRUCT-CAT mutants with *µ* = 200. Values shown here are IC90 of the mutant divided by IC90 of the WT for each parameter combination. Linear correlations are computed between log_10_-transformed parameter values and log_10_-transformed fold-increase in IC90 for each cell type.

